# Plasmid-chromosome transcriptional crosstalk in multidrug resistant clinical enterobacteria

**DOI:** 10.1101/2024.08.08.607126

**Authors:** Laura Toribio-Celestino, Alicia Calvo-Villamañán, Cristina Herencias, Aida Alonso-del Valle, Jorge Sastre-Dominguez, Susana Quesada, Didier Mazel, Eduardo PC Rocha, Ariadna Fernández-Calvet, Alvaro San Millan

**Affiliations:** Centro Nacional de Biotecnología (CNB-CSIC), Madrid, Spain; Servicio de Microbiología, Hospital Universitario Ramón y Cajal and Instituto Ramón y Cajal de Investigación Sanitaria, Madrid, Spain; Centro de Investigación Biológica en Red de Enfermedades Infecciosas, Instituto de Salud Carlos III, Madrid, Spain; Institut Pasteur, Université de Paris Cité, CNRS UMR3525, Bacterial Genome Plasticity, Paris, France; Institut Pasteur, Université de Paris Cité, CNRS UMR3525, Microbial Evolutionary Genomics, Paris, France; Centro de Investigación Biológica en Red de Epidemiología y Salud Pública, Instituto de Salud Carlos III, Madrid, Spain

## Abstract

Conjugative plasmids promote the dissemination and evolution of antimicrobial resistance in bacterial pathogens. However, plasmid acquisition can produce physiological alterations in the bacterial host, leading to potential fitness costs that determine the clinical success of bacteria-plasmid associations. In this study, we used a transcriptomic approach to characterize the interactions between a globally disseminated carbapenem resistance plasmid, pOXA-48, and a diverse collection of multidrug resistant clinical enterobacteria. Although pOXA-48 produced mostly strain-specific transcriptional alterations, it also led to the common overexpression of a small chromosomal operon present in *Klebsiella* spp. and *Citrobacter freundii* strains. This operon included two genes coding for a pirin and an isochorismatase family proteins (*pfp* and *ifp*), and showed evidence of horizontal mobilization across Proteobacteria species. Combining genetic engineering, transcriptomics, and CRISPRi gene silencing, we showed that a pOXA-48-encoded LysR regulator is responsible for the plasmid-chromosome crosstalk. Crucially, the operon overexpression produced a fitness benefit in a pOXA-48-carrying *K. pneumoniae* clinical strain, suggesting that this crosstalk promotes the dissemination of carbapenem resistance in clinical settings.

## Introduction

Antimicrobial resistance (AMR) is becoming a leading global health problem, with AMR infections increasing every year^1,2^. Conjugative plasmids are extrachromosomal genetic elements that can carry diverse accessory genes, including AMR genes, and can transfer them horizontally to nearby cells^3^. Many conjugative plasmids have a broad host range and can thus rapidly disseminate AMR genes across bacterial species^4^. This is especially relevant in clinical settings, where antibiotic pressure persistently selects for AMR plasmid-carrying bacteria^5^. One of the most concerning groups of resistant bacteria are carbapenemase-producing Enterobacteriaceae (CPE)^6^. CPE are able to hydrolyze carbapenems, an important group of antibiotics typically used in hospitals as a last-resort option to treat multidrug resistant (MDR) infections. There are many types of globally disseminated carbapenemases^6^. One relevant example is OXA-48, which is usually encoded in the conjugative plasmid pOXA-48 (plasmid taxonomic unit L/M). OXA-48 was first identified in Turkey in 2001, and is currently involved in hospital outbreaks worldwide^6,7^. Importantly, although pOXA-48 can be carried by many enterobacterial species, it is strongly associated with certain high-risk clones of *Klebsiella pneumoniae* (e.g. of the sequence types ST11 and ST307)^8,9^. Understanding the bacteria-plasmid interactions that shape these successful associations is crucial to controlling AMR dissemination.

When plasmids are acquired by a new bacterial host, they usually disrupt the cell’s transcriptional profile, and this can translate into a fitness cost^10,11^. Chromosomal genes with altered expression are normally related to the metabolism of nucleotides, amino acids and energy sources, translation and transcription, secretion and transport systems, signaling and motility^10,12–21^. These transcriptional changes could just reflect side effects of plasmid carriage or readjustments made to compensate for the new energy requirements and the disruption of cellular homeostasis that result from expressing foreign genes^11,22^. For instance, fitness costs associated with carriage of the pQBR103 or pQBR57 megaplasmids of *Pseudomonas fluorescens* are relieved by mutations in the global regulatory system GacAS, which restore the plasmid-free transcriptional profile^13,23^.

Moreover, there is growing evidence that plasmids may modulate transcriptome changes to manipulate the bacterial host behavior as a strategy to increase their own transmissibility^10,24^. Plasmids have been repeatedly described to reduce the expression of chromosomal genes related to bacterial motility^14,25,26^, presumably to facilitate physical interaction with potential recipients for conjugation^10^. Interestingly, plasmids can employ other strategies to reduce bacterial motility, including translational regulation^27^, or direct protein interactions with the flagella^28^. Other transcriptional modulations could aim at increasing the fitness of the host in new niches to ultimately ensure the plasmid’s vertical transmission^10,24^. For example, plasmid pLL35 induces an increase in the expression of genes involved in anaerobic metabolism in *Escherichia coli*, which could help the host bacterium to colonize the mammalian gut^18^. Moreover, different virulence plasmids have been reported to regulate the expression of chromosomal genes associated with increased virulence^29–32^ or parasitic intracellular growth^33^, promoting a pathogenic lifestyle. All these bacteria-plasmid interactions ultimately shape plasmid distribution and evolution. However, despite the importance of plasmids in the evolution of AMR, we are only starting to understand the transcriptional impact of AMR plasmids on their bacterial hosts, and very few studies have analyzed the effects of relevant AMR plasmids on clinical bacterial strains^15,18,25,34–37^.

In this study, we analyzed the transcriptional response associated with carrying plasmid pOXA-48 in 11 MDR clinical enterobacterial strains of four different species. Our analyses revealed that pOXA-48 produces variable responses on their hosts, but commonly affects processes related to metabolism, transport, cellular organization and motility. More notably, we demonstrated that a pOXA-48-encoded LysR transcriptional regulator directly increases the expression of a small chromosomal operon in *Klebsiella* spp. and *Citrobacter freundii*. This operon showed signs of having been horizontally mobilized across Proteobacteria, suggesting that interactions between mobile genetic elements may shape the evolution of plasmid-mediated AMR in clinical enterobacteria. Moreover, our results suggest that the plasmid-chromosome crosstalk produces a fitness benefit in pOXA-48-carrying clinical strains.

## Results

### Transcriptomic analysis of pOXA-48-carrying clinical enterobacteria

To study the transcriptional impact of pOXA-48 carriage in clinically relevant bacteria, we analyzed 11 MDR enterobacterial strains isolated from the gut microbiota of patients admitted to a large University Hospital in Madrid, Spain (Supplementary Table 1 and Supplementary Fig. 1). These 11 strains belong to four different species (*K. pneumoniae,* n = 7; *E. coli,* n = 2; *K. variicola*, n = 1; *C. freundii*, n = 1) and are representative of the phylogenetic diversity of MDR enterobacteria isolated in this hospital (Supplementary Fig. 1A)^9^. We also included in our analyses the laboratory-adapted *E. coli* MG1655 strain. As observed in our previous studies on the distribution of fitness effects of pOXA-48 in clinical enterobacteria, pOXA-48 produced a wide range of fitness effects in these strains (Supplementary Fig. 1B)^38,39^. For each strain, we had previously obtained pOXA-48-free and pOXA-48-carrying isogenic clones by curing (using CRISPR-Cas9 technology) or introducing (by conjugation) this plasmid^38–40^. To perform the differential expression (DE) analysis, we sequenced the transcriptomes of all clones (pOXA-48-carrying and pOXA-48-free, n = 24 clones, 2-4 biological replicates per clone, see Methods).

### Conserved expression of pOXA-48 genes across strains

Although the structure and sequence of plasmid pOXA-48 is quite conserved across clinical enterobacteria, we have previously described multiple pOXA-48 variants carrying different indels and/or SNPs^41^. Therefore, we first performed variant calling on the transcriptomic data and confirmed that the polymorphisms identified in pOXA-48 did not alter expression of the affected genes (Fig. 1 and Supplementary Tables 2 and 3). We identified SNPs upstream of the *repA* gene of pOXA-48 in all replicates of *K. pneumoniae* KPN07 and KPN10 and in one replicate of EC10 (Supplementary Table 2). These mutations, which increase pOXA-48 copy number (PCN), have been previously described in clinical strains^41^, so we decided to maintain them in the subsequent analyses.

**Fig. 1.**
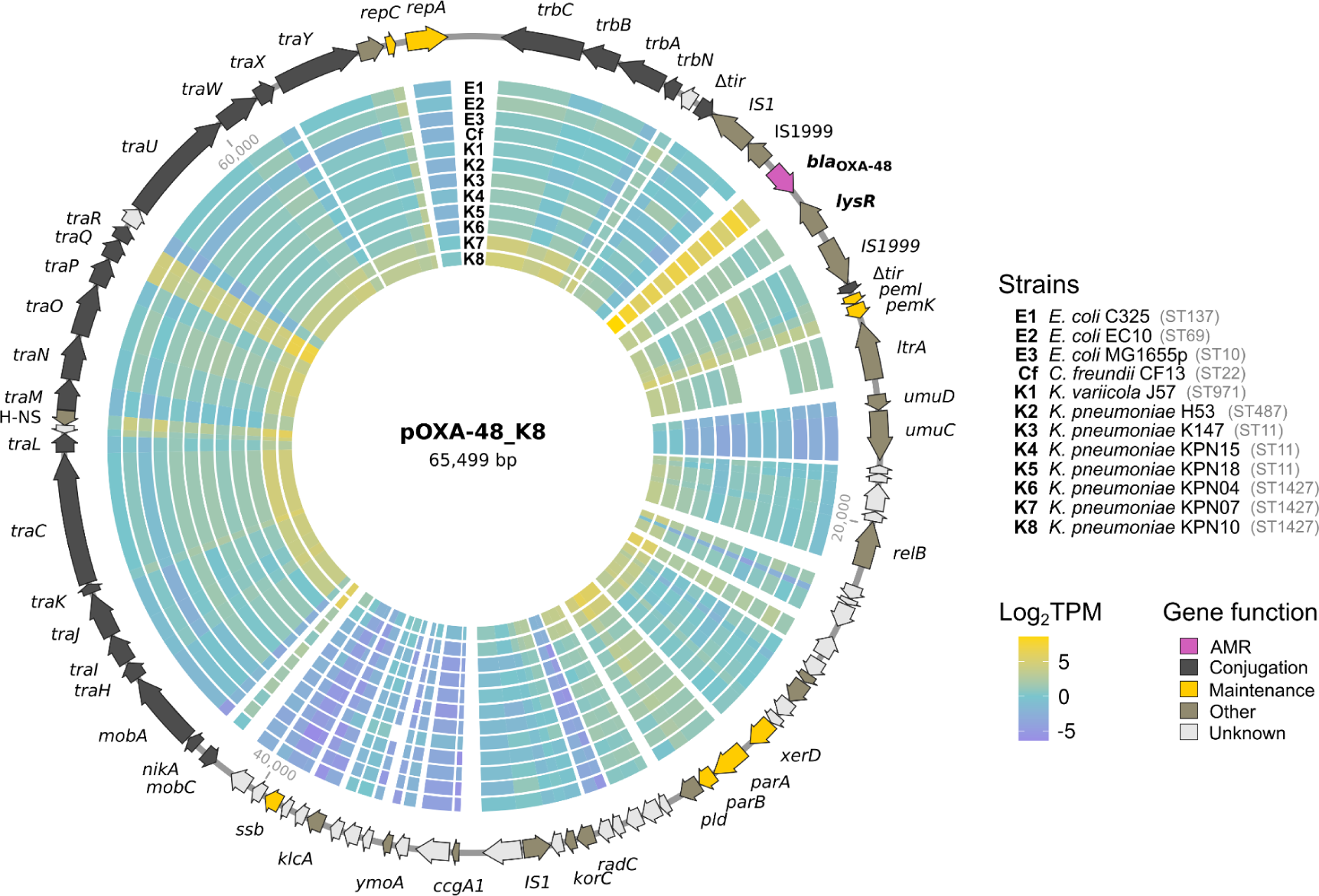
Expression of pOXA-48 genes. The genetic map of pOXA-48 shows the direction of transcription of genes as arrows, with colors indicating gene function. Each concentric circle of the heatmap represents the expression of pOXA-48 genes of each strain, measured as the median log_2_TPM (Transcripts Per Million) between replicates, normalized by median chromosomal TPM values of the corresponding strain. Log_2_TPM > 0 indicates that pOXA-48 genes are expressed more than the median expression of chromosomal genes, while log_2_TPM < 0 indicates that pOXA-48 gene expression is below the median chromosomal TPM values. Gaps in the heatmap mean that the pOXA-48 sequence of the reference genome does not include the gene or the gene was not annotated (see Supplementary Table 2).

Next, we analyzed the expression of pOXA-48 genes in the different genetic backgrounds of the 12 enterobacterial strains. Gene expression was calculated as Transcripts Per Million (TPM). To facilitate comparison between strains, TPM values were normalized by the median chromosomal TPM values of the corresponding strain. All plasmid genes were transcribed (Fig. 1, Supplementary Fig. 2 and Supplementary Table 3), in contrast to other AMR plasmids, where up to 40% of genes were silent^15,42,43^. We did not observe distinctive transcriptional profiles of pOXA-48 associated with the different host genetic backgrounds. However, strains KPN07 and KPN10 and the replicate 3 of EC10 showed overall higher pOXA-48 expression (Fig. 1, Supplementary Fig. 2 and Supplementary Table 3), in agreement with the increased PCN caused by the mutation upstream *repA*. The most expressed gene was the carbapenemase *bla*_OXA-48_ (Fig. 1, Supplementary Fig. 2 and Supplementary Table 3). Other works have also reported AMR genes to be amongst the most expressed plasmid genes^14,15,26,42,44^. Other highly transcribed genes were *traQ* and *traR*, involved in pilin formation and plasmid transfer, an H-NS-like regulatory protein-coding gene, and the toxin and antitoxin *pemK* and *pemI* genes, associated with plasmid maintenance through post segregational killing (Fig. 1 and Supplementary Table 3). The analysis of expression of other resident plasmid’s genes is described in Supplementary Information.

### Strain-specific transcriptomic changes associated with pOXA-48 carriage

pOXA-48 produced variable, strain-specific transcriptomic changes on the chromosome of its enterobacterial hosts (Supplementary Fig. 3), ranging from only six significant differentially expressed genes (DEGs) in *K. pneumoniae* H53 to more than a thousand DEGs in the strains that carried a pOXA-48 variant with increased PCN (EC10, KPN07 and KPN10, 23-37% of total chromosomal genes) (Supplementary Table 4). In the remaining clinical strains, pOXA-48 induced the DE of 2-15% of chromosomal genes (Supplementary Table 4). Several studies have also reported strain-specific transcriptional changes associated with plasmid carriage^14,18,45^. Interestingly, only 63 genes were DE in the laboratory *E. coli* MG1655 strain (1.4% of total chromosomal genes) (Supplementary Table 4), which is surprising because pOXA-48 caused the highest fitness costs on this strain (Supplementary Fig. 1). However, in line with other studies^18,20^, the number of DEGs did not correlate with the fitness cost imposed by pOXA-48 (Spearman’s rank correlation, *ρ* = -0.1*, P* = 0.75, Supplementary Fig. 4).

To find commonly enriched functions between strains, we performed gene set enrichment analysis (GSEA) on the lists of raw DEGs with Gene Ontology (GO) functional annotations (Fig. 2, Supplementary Figs. 5 and 6 and Supplementary Table 5). We also counted the number of significant up- and downregulated genes for each GO Biological Process across strains (Supplementary Table 6). pOXA-48 carriage produced the most significant impact on host metabolism (Fig. 2, Supplementary Fig. 5 and Supplementary Tables 5 and 6). The affected metabolic pathways and the extent of the changes varied across strains, but carbohydrate metabolism (including glycolysis and the TCA and glyoxylate cycles) and respiration (both aerobic and anaerobic, as well as the electron transport chain) were altered in multiple strains (Fig. 2 and Supplementary Tables 5 and 6). This suggests that pOXA-48 could potentiate the exploration of alternative energy sources in its hosts, as seen for other plasmids^18,20,21^. For example, nitrate assimilation and metabolism were upregulated in pOXA-48-carrying KPN07, EC10 and KPN10 (Fig. 2, Supplementary Fig. 5 and Supplementary Tables 4, 5 and 6).

**Fig. 2.**
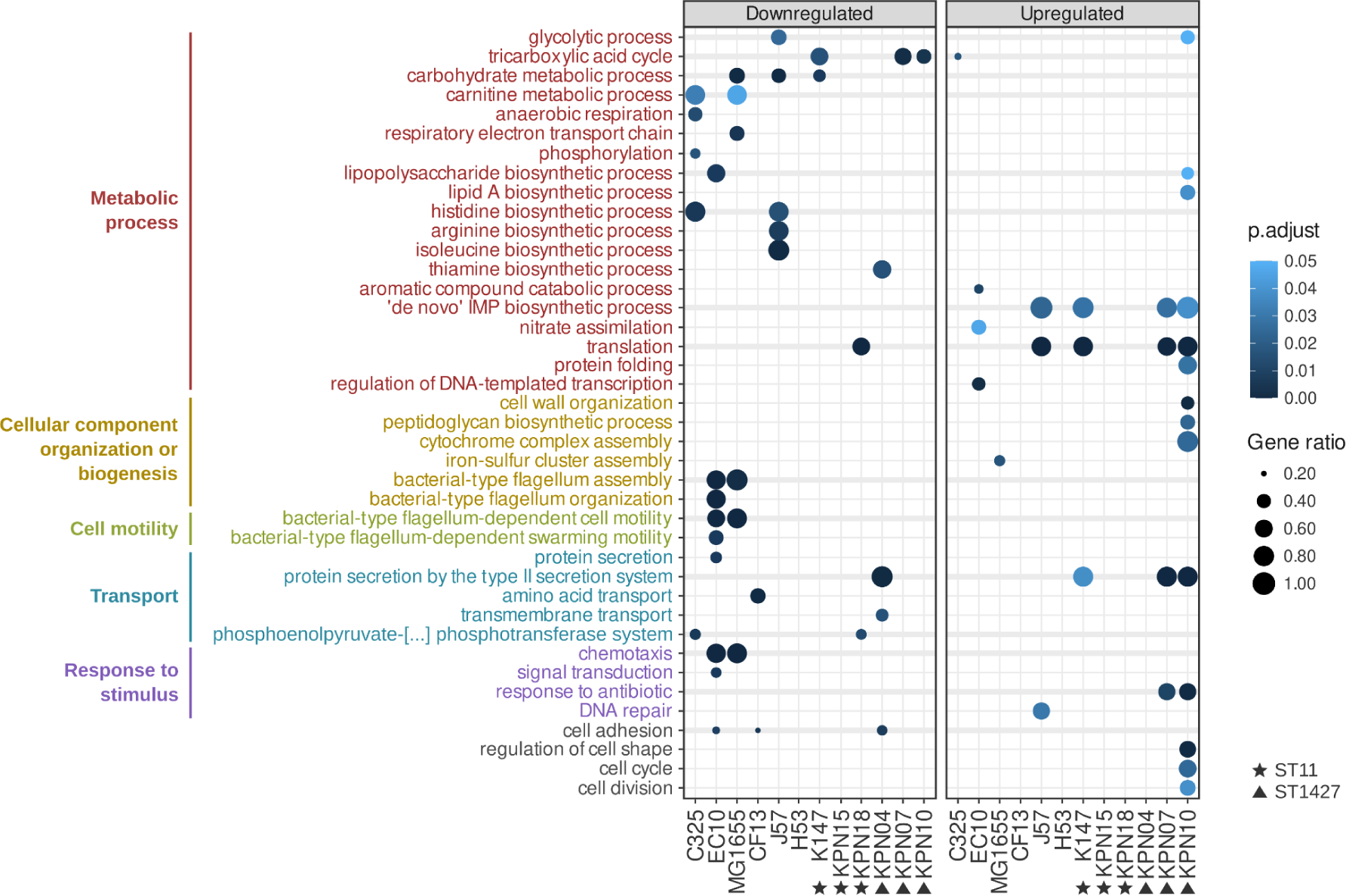
Transcriptomic responses to pOXA-48 carriage. Biological Processes (BP) enriched in pOXA-48-carrying enterobacteria. Gene set enrichment analysis (GSEA) was performed on the lists of raw DEGs annotated with Gene Ontology terms for BP (see Methods). Downregulated and upregulated enriched BP are separated in two panels, and are represented by circles. Thicker horizontal lines indicate BP enriched in more than one strain. The size of the circles indicate the ratio of the number of enriched genes of a specific BP by the number of total genes annotated for that BP. The adjusted *p* value for each enriched BP and strain is represented in a color gradient. Strains belonging to the same Sequence Type (ST) are indicated with shapes.

pOXA-48 also produced an overall impact in bacterial motility. Genes involved in flagella-dependent motility were strongly downregulated in 2 of the 3 *E. coli* strains: EC10 and MG1655 (Fig. 2, Supplementary Fig. 6 and Supplementary Tables 4, 5 and 6). Cell adhesion mediated by fimbriae tended to be downregulated in several strains, especially in *E. coli* EC10, *C. freundii* CF13 and *K. pneumoniae* KPN04 (Fig. 2, Supplementary Fig. 6 Supplementary Tables 4, 5 and 6). Chemotaxis was also downregulated in EC10 and MG1655 (Fig. 2 and Supplementary Tables 4, 5 and 6). Reduced motility is a common plasmid effect, which could facilitate its horizontal transmission through conjugation^10,14,25,26^.

### pOXA-48 induces the overexpression of a small and mobile chromosomal operon

Very few common genes were significantly DE across the strains, and those who were showed different directions of DE (Supplementary Fig. 3C and Supplementary Table 4). However, we observed a tuned overexpression of two adjacent chromosomal genes (*pfp* and *ifp*) coding for a pirin and isochorismatase family proteins in all *Klebsiella* spp. strains and in *C. freundii* (Fig. 3A and Supplementary Table 4). Both genes were predicted to form a small operon (with a 97% probability as reported by Operon-mapper). A LysR family transcriptional regulator was encoded upstream of the operon, in antisense, and could therefore regulate its expression (Fig. 3A). From here on, we will refer to these three genes as the *lysR-pfp-ifp* cluster. Crucially, none of the *E. coli* strains encoded the *lysR-pfp-ifp* cluster (Fig. 3A).

**Fig. 3.**
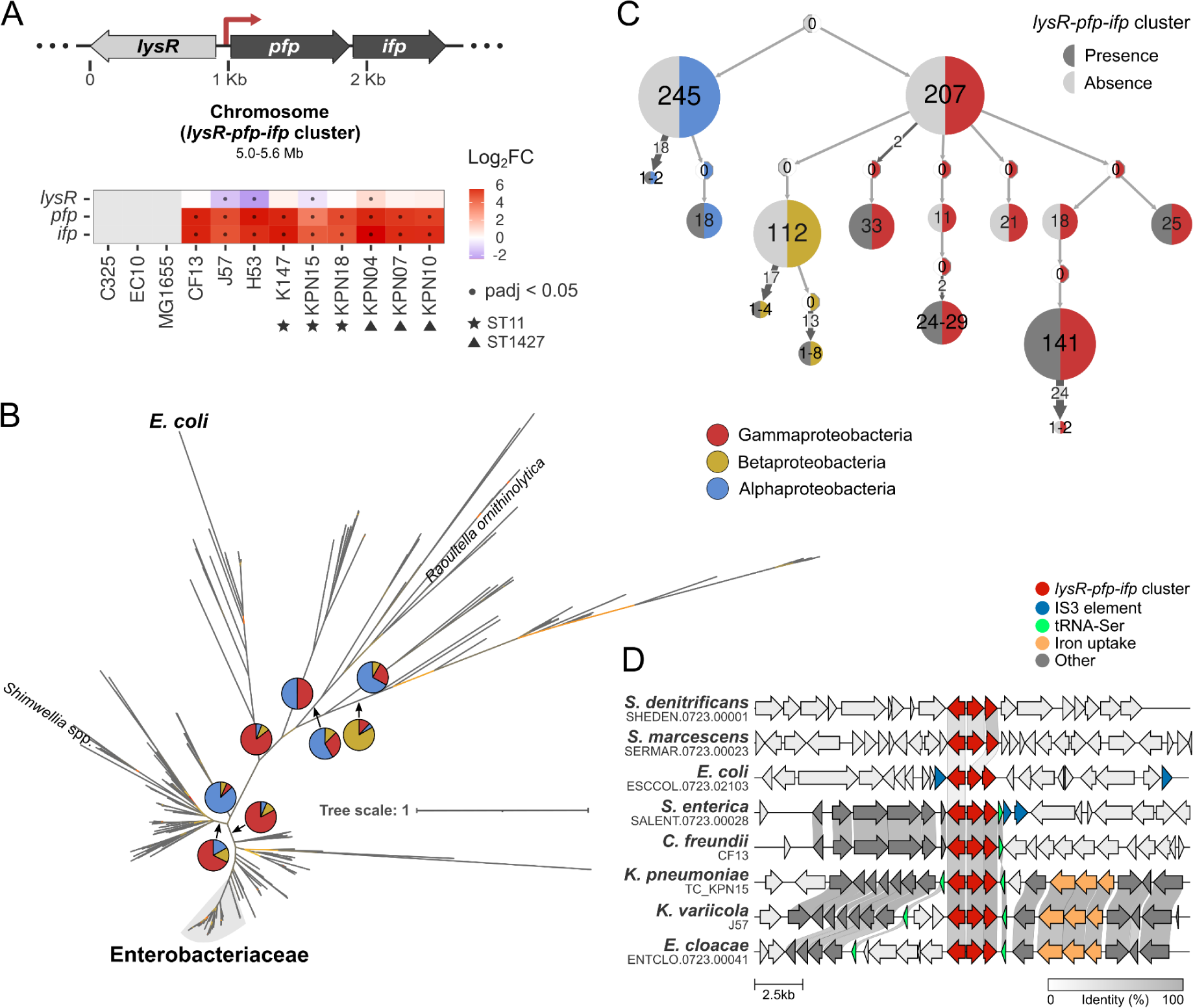
Overexpression of a small, horizontally-acquired, chromosomal operon. (A) Schematic representation (top figure) and heatmap of the differential expression (log2 fold change [FC], bottom figure) of a small chromosomal operon, comprised of the *pfp* and *ifp* genes, and the *lysR* transcriptional regulator gene that could control its expression (*lysR-pfp-ifp* cluster). Differential expression is performed by comparing the pOXA-48-carrying strains to their respective pOXA-48-free counterparts. *E. coli* (C325, EC10 and MG1655) does not encode the *lysR-pfp-ifp* cluster, thus heatmap tiles are colored in gray. Significant log2FC are indicated by a dot. Strains belonging to the same Sequence Type (ST) are indicated with shapes. (B) Unrooted phylogenetic tree of the concatenated alignments of the protein sequences of the *lysR-pfp-ifp* cluster genes in Proteobacteria (n = 698) (see Methods). Pie charts show the proportion of Proteobacteria strains belonging to the Gamma-(red), Beta-(yellow) or Alphaproteobacteria (blue) classes included below each of the indicated internal branches. Enterobacteriaceae strains that branch from the same node are shaded in gray; other Enterobacteriaceae species not branching from that node are indicated. Bootstrap values are represented with a color gradient in the branches of the tree: low (≥8), medium and high (≤100) bootstrap values are colored in red, yellow and gray, respectively. Tree scale represents the average number of substitutions per site. (C) Ancestral character reconstruction of the *lysR-pfp-ifp* cluster in Proteobacteria (n = 1,686; see Methods). The Proteobacteria phylogenetic tree is compressed vertically and horizontally based on the Proteobacteria class (right halves of circles) and the presence/absence of the *lysR-pfp-ifp* cluster (dark and light gray, respectively; left halves of circles). White circle halves indicate that the state of the respective character could not be inferred, which occurs in the internal nodes of the tree, indicated with 0s. Numbers inside circles indicate the number of tree tips that possess the corresponding characters after vertical compression. Numbers in arrows indicate the number of identical subtrees horizontally merged. (D) Genomic neighborhood of the *lysR-pfp-ifp* cluster in a representative subset of Proteobacteria strains: *Shewanella denitrificans*, *Serratia marcescens*, *E. coli*, *Salmonella enterica*, *C. freundii*, *K. pneumoniae*, *K. variicola* and *Enterobacter cloacae*. Genes are color-coded as indicated at the top. Homologous genes between strains and their percentage of sequence identity are indicated with a gray shading.

A search on the STRING database revealed that the *lysR-pfp-ifp* cluster is present in many, but not all, species of Proteobacteria (Supplementary Fig. 7). This particular distribution led us to investigate the possible origin of the *lysR-pfp-ifp* cluster. For this, we used the RefSeq database of all non-redundant complete genomes of Proteobacteria (n = 22,631) and identified the *lysR-pfp-ifp* cluster in 5,983 Gamma-(18.6% of species), 191 Beta-(18.9% of species) and 172 Alphaproteobacteria (8.8% of species) genomes (Supplementary Table 7; see Methods). Next, we constructed a phylogenetic tree of 698 representative *lysR-pfp-ifp* cluster-encoding strains of Proteobacteria from the concatenated alignments of the LysR, PFP and IFP protein sequences (Fig. 3B). We also constructed a separate tree for each of the proteins (Supplementary Figs. 8, 9 and 10), which showed similar tree topologies, suggesting the three genes are evolutionarily related. Curiously, the tree of the *lysR-pfp-ifp* cluster did not branch according to the taxonomic class of the strains, but rather all clades included a mix of classes of Proteobacteria (Fig. 3B), suggesting the *lysR-pfp-ifp* cluster experienced frequent horizontal transfer. This was supported by ancestral character reconstruction, where the *lysR-pfp-ifp* cluster appeared at the tips of the tree and outer internal nodes in all classes of Proteobacteria, and was often absent in more internal nodes, suggesting horizontal acquisition (Fig. 3C and Supplementary Fig. 11).

In line with this hypothesis, the genes neighboring the *lysR-pfp-ifp* cluster differed between species of Proteobacteria, suggesting it could have been horizontally acquired and integrated at different genomic regions (Fig. 3D and Supplementary Fig. 12). Accordingly, of the 3,105 *E. coli* strains in the database, only two encoded a *lysR-pfp-ifp* cluster, which was located near genes annotated as transposases of the IS3 family (Fig. 3B and 3D). The *lysR-pfp-ifp* cluster was also inserted between homologous tRNAs in different species (Fig. 3D), which are known integration sites for mobile genetic elements ^46^. Moreover, the *lysR-pfp-ifp* cluster showed different GC content (difference ranging from 0.4% to 9.2%, median 2.8%; Supplementary Table 8) and lower codon adaptation index compared to the host bacteria chromosomes for different species (median CAI between of *lysR* = 0.42, *pfp* = 0.57 and *ifp* = 0.6, median CAI of ribosomal genes = 0.76; Supplementary Table 8), further supporting the hypothesis of high horizontal mobility of the *lysR-pfp-ifp* cluster.

### A pOXA-48-encoded LysR regulator increases the expression of the *pfp-ifp* operon

pOXA-48 encodes a LysR transcriptional regulator downstream the *bla*_OXA-48_ gene (Fig. 4A, see Supplementary Information for a detailed analysis of LysR_pOXA-48_ sequence conservation across pOXA-48 plasmids). Curiously, LysR_pOXA-48_ shows the highest protein sequence identity (34.2% in *C. freundii* and 36.7-37% in *Klebsiella* spp.) with the LysR that putatively regulates the expression of the *pfp-ifp* operon in comparison to other chromosomal LysRs (Supplementary Table 9). Thus, we hypothesized that LysR_pOXA-48_ could be activating the expression of the operon genes (Fig. 4A).

**Fig. 4.**
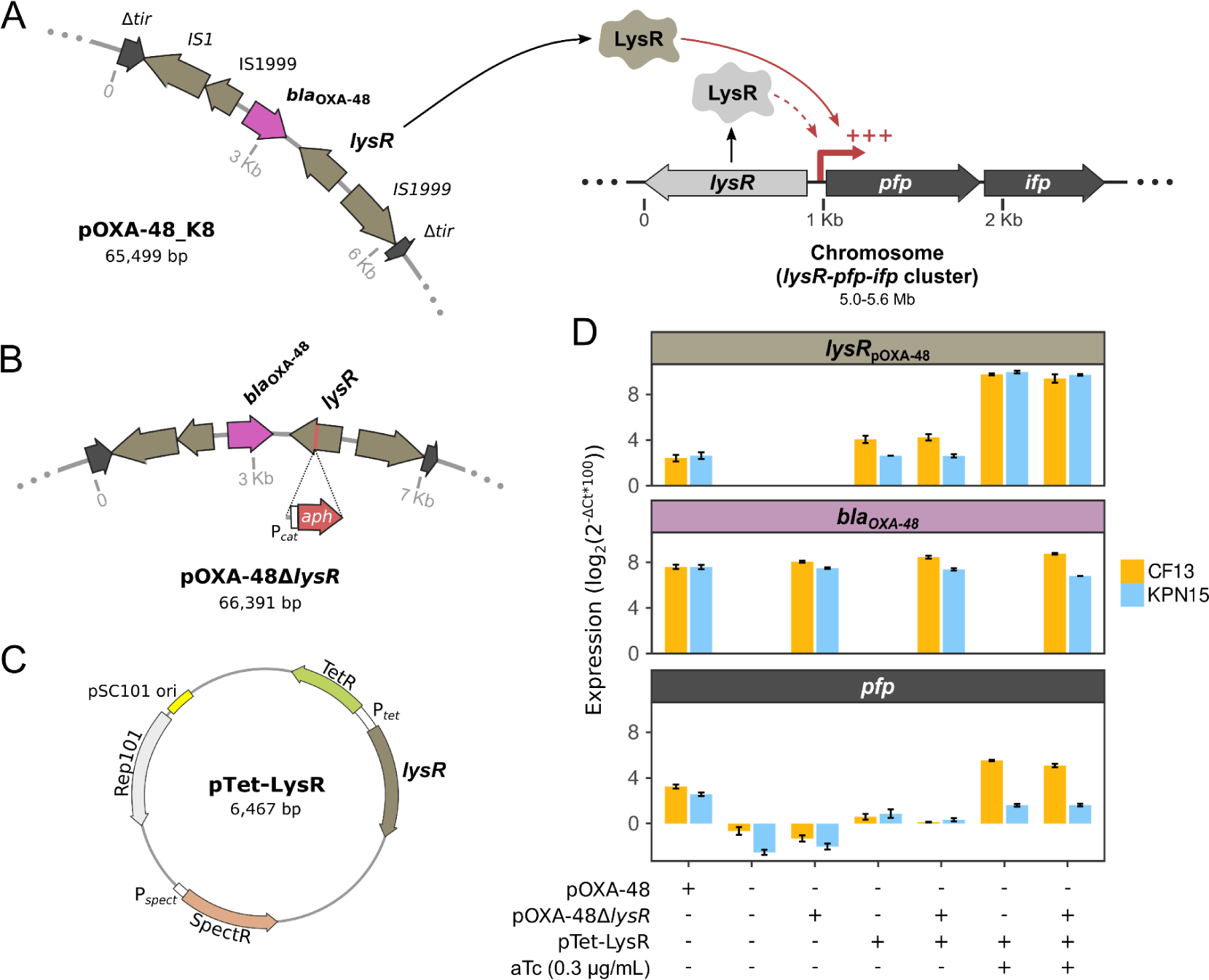
Plasmid-chromosome transcriptional crosstalk. (A) Schematic representation of the plasmid-chromosome crosstalk mediated by the pOXA-48-encoded LysR transcriptional regulator. In the presence of pOXA-48, the overexpression of the operon genes is mediated by LysR_pOXA-48_ (solid red arrow). The thick red arrow represents the putative promoter of the operon. Chromosome and plasmid are represented at different scales, indicated in the figure. (B) Zoom in on the pOXA-48Δ*lysR* plasmid. The *lysR* gene of pOXA-48 was knocked-out by inserting the *aph(3’)-IIa* resistance gene. (C) Map of the pTet-LysR plasmid. The expression of *lysR*_pOXA-48_ is controlled by the inducible promoter P*_tet_*. (D) RT-qPCR results for the expression of *lysR*_pOXA-48_, *bla*_OXA-48_ and the *pfp-ifp* operon (*pfp* levels as proxy for the entire operon). Levels of expression are normalized to the endogenous control *rpoB* and represented as the log_2_FC of gene expression compared to *rpoB.* Error bars represent the standard error of the mean (n = 2 biological replicates, each as the median of 3 technical replicates).

To test this hypothesis, we constructed a version of pOXA-48 with *lysR*_pOXA-48_ interrupted by a kanamycin-resistance cassette (pOXA-48Δ*lysR,* Fig. 4B) and conjugated it into two pOXA-48-free clinical strains: *K. pneumoniae* KPN15 and *C. freundii* CF13. No *E. coli* strain was included in this part of the study due to the absence of the *lysR-pfp-ifp* cluster in all of our *E. coli* strains. Using RT-qPCR, we compared the levels of expression of the *pfp-ifp* operon in the presence of pOXA-48 or pOXA-48Δ*lysR,* and in the absence of either plasmid (Fig. 4D). Our results confirmed that the *pfp-ifp* operon is overexpressed (ca. 20-fold) in the presence of pOXA-48 (Dunnett’s test *P* = 0.0008 for KPN15, *P* = 0.034 for CF13), but not in the presence of pOXA-48Δ*lysR* (Dunnett’s test *P* = 0.3 for KPN15 and *P* = 0.3 for CF13), compared to the plasmid-free strains, both for KPN15 and CF13 (Fig. 4D).

To complement the deletion of *lysR,* we constructed the expression vector pTet-LysR (Fig. 4C). This vector expresses *lysR*_pOXA-48_ under the pTet promoter, inducible by anhydrotetracycline (aTc). The LysR_pOXA-48_ expressed from the induced pTet-LysR vector led to the overexpression of the *pfp-ifp* operon, showing that LysR_pOXA-48_ is necessary and sufficient for the transcriptional activation of the *pfp-ifp* operon (Fig. 4D). Additionally, our RT-qPCR results revealed that LysR_pOXA-48_ plays no role in the expression of *bla*_OXA-48_, as levels of *bla*_OXA-48_ were similar using pOXA-48 or pOXA-48Δ*lysR* (*t*-test *t*(1.19) = 0.49, *P* = 0.7 for KPN15, *t*(1.45) = -2.16, *P* = 0.21 for CF13; Fig. 4D).

### The LysR-mediated transcriptional crosstalk is specific to the operon genes

To further characterize the regulatory functions of LysR_pOXA-48_, we sequenced the transcriptomes of *K. pneumoniae* KPN15 and *C. freundii* CF13, carrying pOXA-48 or pOXA-48Δ*lysR*, and performed DE analysis. When analyzing the expression of pOXA-48 genes, none was DE in the same direction in both strains –including *bla*_OXA-48_, in line with our RT-qPCR results–, and the magnitude of DE was moderate (log_2_FC ranging from -0.56 to 1.46, Supplementary Table 10). This indicates that LysR_pOXA-48_ does not regulate the expression of any pOXA-48 gene.

The operon genes *pfp* and *ifp* were overexpressed when comparing the pOXA-48 carriers against the LysR_pOXA-48_-deficient strains carrying pOXA-48Δ*lysR*, further confirming that LysR_pOXA-48_ is responsible for the increased expression of the operon (mean log_2_FC of *pfp* = 5.54 and *ifp* = 4.95; Fig. 5 and Supplementary Table 10). Apart from the operon genes, few other chromosomal genes were strongly DE (log_2_FC ranging from -2.53 to 2.67, mean = 0.007) or were DE in the same direction in KPN15 and CF13 (Supplementary Table 10), suggesting that LysR_pOXA-48_ only activates the expression of the operon. To investigate this further, we analyzed the DE of genes that could be under the control of other LysR regulators. We considered a gene to be putatively controlled by a LysR if a *lysR* gene was located upstream and in antisense of the gene, as in the *lysR-pfp-ifp* cluster (Fig. 5). Only five genes putatively controlled by a LysR (excluding *pfp*) were significantly DE in CF13 and KPN15 (Fig. 5). However, they were only marginally DE (log_2_FC ranging from -1.4 to 0.37) and none of them were DE in the same direction across strains when comparing pOXA-48 carriers against plasmid-free strains (Supplementary Table 4). These results suggest that the LysR-mediated plasmid-chromosome crosstalk is specific to the operon genes.

**Fig. 5.**
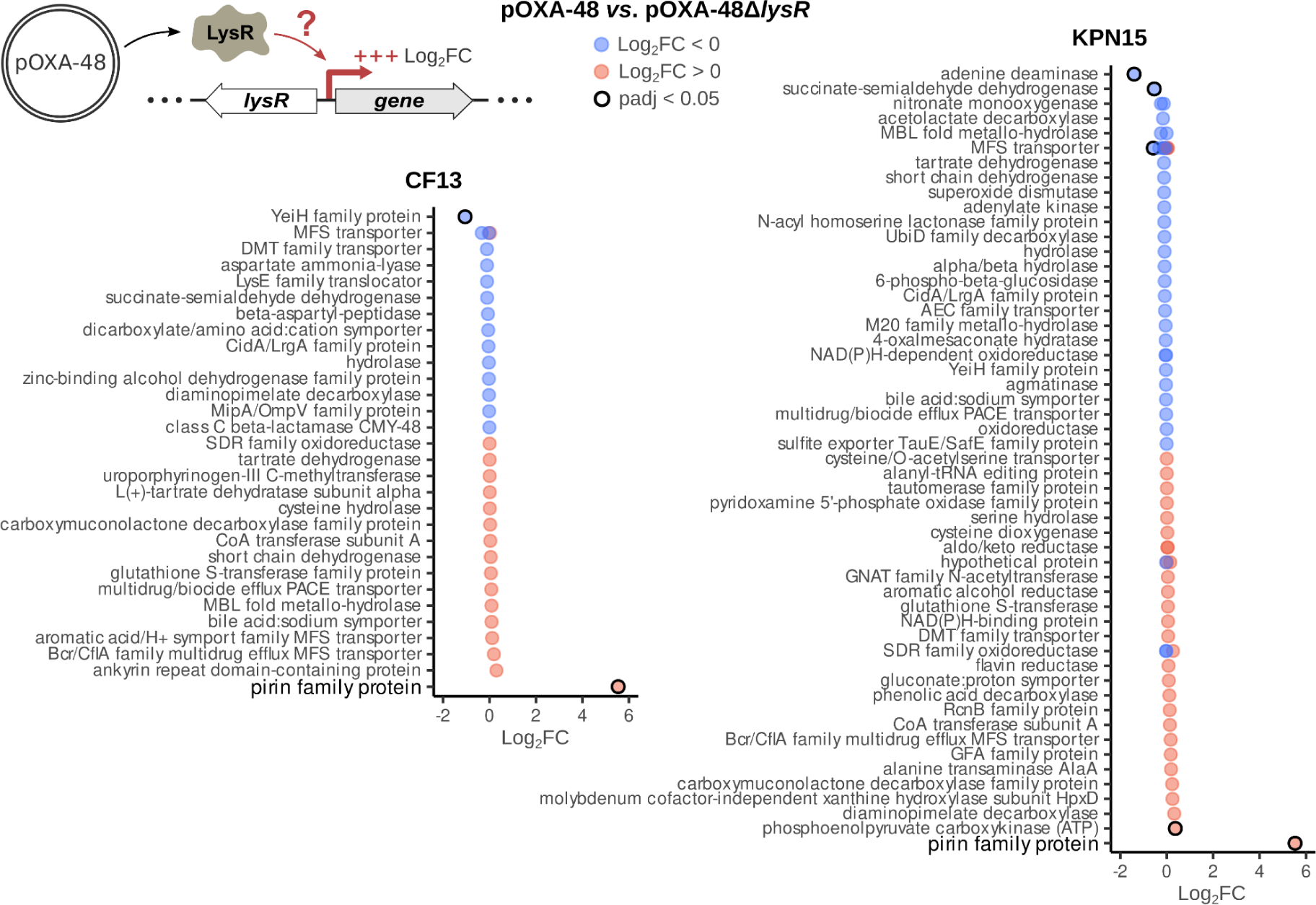
Differential expression of chromosomal genes putatively controlled by other LysRs. Genes were considered to be potentially regulated by a LysR transcriptional regulator if a *lysR* gene was preceding the gene in antisense (top schematic representation). The log_2_FoldChanges (comparing pOXA-48-carrying to pOXA-48Δ*lysR-*carrying counterparts) of the genes (represented as gene products) putatively controlled by LysRs are plotted for CF13 (left) and KPN15 (right). The pirin family protein of the *lysR-pfp-ifp* cluster is emphasized. Red and blue circles represent up- and downregulation, respectively. Outlined circles indicate significant log_2_FC (adjusted *p* value < 0.05).

### Functional analysis of the operon genes

Next, we decided to study the function of the *pfp-ifp* operon genes. Many members of the pirin family protein possess quercetinase activity, oxidizing the flavonoid quercetin^47^. To test the quercetinase activity of the PFP protein, we cultured *C. freundii* CF13 and *K. pneumoniae* KPN15 carrying either pOXA-48 or pOXA-48Δ*lysR* in different concentrations of quercetin solubilized in DMSO (Fig. 6A and Supplementary Fig. 13). Quercetin possesses antimicrobial activity^48^, thus we hypothesized that if PFP degrades quercetin, strains carrying pOXA-48Δ*lysR* –producing less PFP– would present a growth defect compared to pOXA-48-carrying strains. By constructing a generalized additive model per strain (GAM; see Methods), we showed that, in KPN15, pOXA-48Δ*lysR*-carrying strains are more susceptible to quercetin than pOXA-48-carrying strains (Fig. 6A, Supplementary Fig. 14 and Supplementary Table 11), which strongly suggests that PFP has quercetinase activity in *K. pneumoniae*. In CF13, on the other hand, we did not observe a clear difference between pOXA-48- and pOXA-48Δ*lysR*-carrying strains (Fig. 6A, Supplementary Fig. 14 and Supplementary Table 11).

**Fig. 6.**
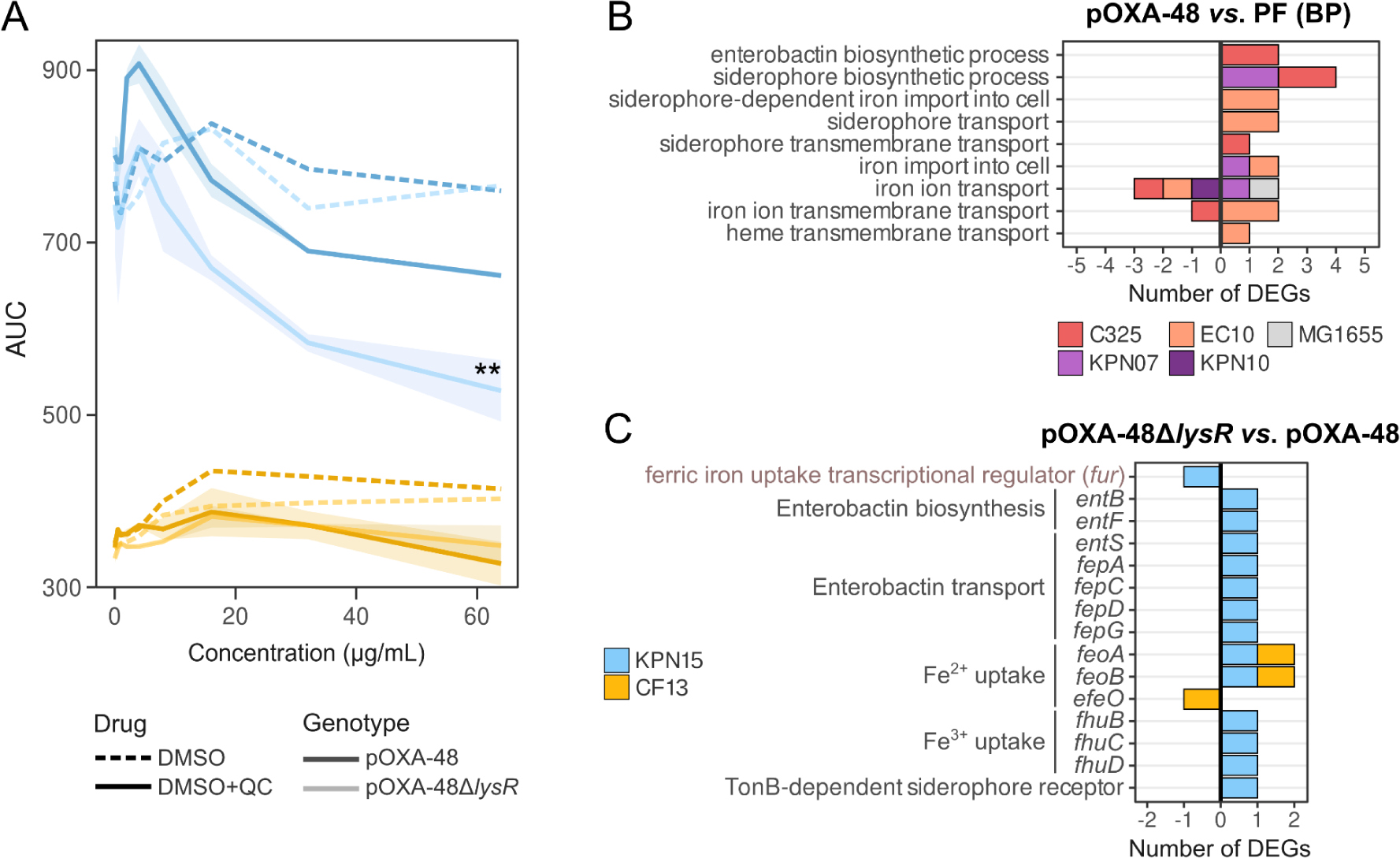
Functional analysis of the operon genes. (A) AUC of growth curves under increasing concentrations of DMSO or DMSO + quercetin (QC) of KPN15 and CF13 carrying pOXA-48 or pOXA-48Δ*lysR*. This data was used to construct a GAM per strain (Supplementary Fig. 14; see Methods). Shading indicates the standard error of the mean (n = 1 for DMSO and n = 2 for DMSO+QC). Asterisks represent statistical significance: * *P* ≤ 0.05, ** *P* ≤ 0.01, *** *P* ≤ 0.001. (B) Number of significant DEGs annotated with biological processes (BP) related to iron transport (comparing pOXA-48-carrying against pOXA-48-free [PF] strains of the first RNA-Seq dataset). Note that genes annotated with more than one of the represented BP will be counted once in each BP. (C) Number of significant DEGs involved in iron transport (comparing pOXA-48Δ*lysR-*carrying against pOXA-48-carrying strains of the second RNA-Seq dataset). In B and C, values >0 represent the count of upregulated genes, and values <0 are downregulated genes.

Members of the isochorismatase-like superfamily, like the EntB enzyme involved in enterobactin biosynthesis, typically hydrolyze the isochorismate to 2,3-dihydroxybenzoate and pyruvate. Enterobactin is a high-affinity bacterial siderophore involved in iron uptake^49^. Interestingly, when comparing pOXA-48-carrying to -free strains, we observed a tendency to overexpress genes related to iron uptake in the *E. coli* strains –that lack the operon– and *K. pneumoniae* KPN07 and KPN10 clinical strains –with higher PCN– (Fig. 6B and Supplementary Fig. 15), suggesting an increased pOXA-48-dependent iron demand in these strains. This tendency was also observed in KPN15 and CF13 carrying the pOXA-48Δ*lysR* variant compared to the strains carrying pOXA-48 (Fig. 6C). Moreover, in this last comparison, KPN15/pOXA-48Δ*lysR* also downregulated the expression of *fur*, a global ferric uptake transcriptional repressor (Fig. 6C). Thus, we hypothesized that IFP could be related to the enterobactin biosynthetic pathway, increasing enterobactin-mediated iron uptake and compensating an increased iron demand caused by pOXA-48 carriage. To determine whether overexpression of the *pfp-ifp* operon increases iron uptake, we measured growth of CF13 and KPN15 under increasing concentrations of the iron chelator bipyridine (Supplementary Fig. 16). However, we did not observe any growth difference in strains carrying pOXA-48 or pOXA-48Δ*lysR* in iron-deficient conditions (Supplementary Fig. 16). The regulation of iron transport is complex, with many different systems involved in iron acquisition. It is therefore possible that the observed adjustments in expression of genes involved in iron import (Fig. 6B-C) could be compensating for the underexpression of the *pfp-ifp* operon, increasing iron acquisition in the pOXA-48Δ*lysR*-carrying strains.

### Overexpression of the *pfp-ifp* operon increases the fitness of a pOXA-48-carrying strain

To investigate the biological relevance of the *pfp-ifp* operon, we first measured the fitness effects associated with *pfp-ifp* overexpression in pOXA-48-free strains. We performed growth curves for both *C. freundii* CF13 and *K. pneumoniae* KPN15 carrying the pTet-LysR vector, which induction leads to *pfp-ifp* expression (Fig. 7A). As a proxy for fitness, we measured the area under the growth curve (AUC) at different levels of aTc induction (note that even with no aTc there is a leaky expression from pTet-LysR, see Fig. 4D). In CF13, aTc produced a moderate growth arrest (one-way ANOVA, CF13: *F*(3,16) = 22.45, *P* < 0.001; KPN15: *F*(3,16) = 0.042, *P* = 0.99), which complicated the interpretation of the results in this strain. Overall, we observed that pTet-LysR produced a beneficial effect in CF13 and a detrimental effect in KPN15, across different aTc concentrations (two-way ANOVA for the interaction between aTc concentration and pTet-LysR, CF13: *F*(3,32) = 7.60, *P* < 0.001; KPN15: *F*(3,32) = 4.31, *P* = 0.014; for Tukey’s multiple comparisons tests see Supplementary Table 11).

**Fig. 7.**
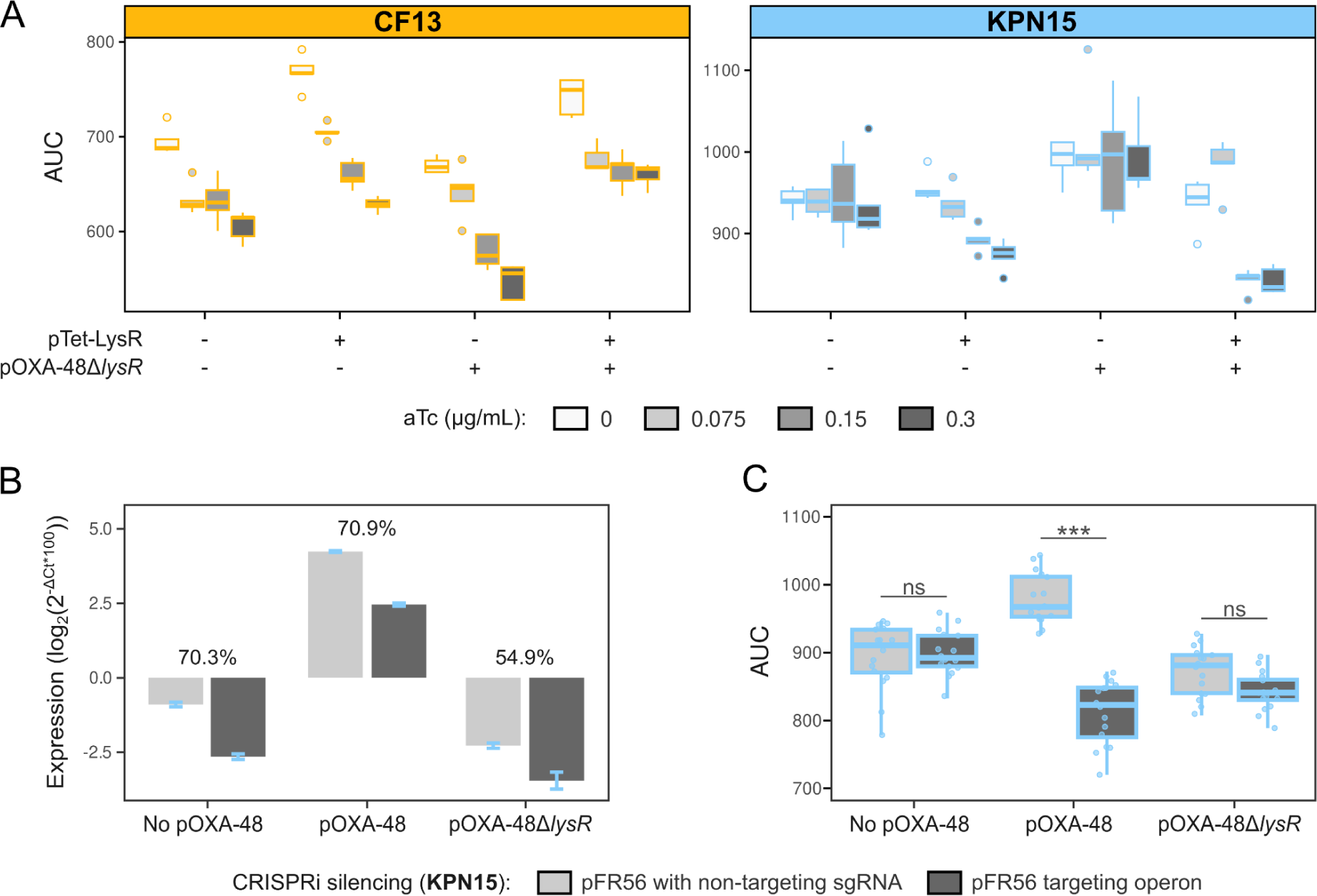
Analysis of the overexpression and repression of the *pfp-ifp* operon. (A) Effect of the overexpression of LysR_pOXA-48_ on the growth of CF13 and KPN15. AUC of plasmid-free and pOXA-48Δ*lysR*-carrying CF13 and KPN15, with expression of LysR_pOXA-48_ in *trans* from the pTet-LysR (n = 5 replicates for each strain/condition pairing). Experiment performed with increasing concentrations of inducer (aTc). (B) Effect of CRISPRi repression on the expression of the *pfp-ifp* operon. RT-qPCR results for *pfp* as proxy of the whole operon, normalized to the endogenous control *rpoB*. Percentages indicate the level of repression of the operon (expression levels of *pfp* in cells with non-targeting pFR56apm *vs*. in cells carrying pFR56apm targeting the *pfp-ifp* operon). Error bars represent the standard error of the mean (n = 2 biological replicates, each as the median of 3 technical replicates). (C) Effect of CRISPRi repression on the growth of plasmid-free, pOXA-48- and pOXA-48Δ*lysR*-carrying KPN15. In boxplots, horizontal lines inside boxes indicate median values, upper and lower hinges correspond to the 25th and 75th percentiles, and whiskers extend to observations within the 1.5x the interquartile range. Individual points show independent replicates (n = 16). Asterisks represent statistical significance: * *P* ≤ 0.05, ** *P* ≤ 0.01, *** *P* ≤ 0.001.

To directly study the effect of *pfp-ifp* overexpression in the pOXA-48-carrying strains, we constructed a CRISPRi system able to silence the *pfp-ifp* operon. The pFR56apm vector carries *Streptococcus pyogenes’* dCas9 under the control of a pHlf promoter, and is easily reprogrammable to target any desired DNA *locus* by changing an sgRNA which is constitutively expressed. To silence the *pfp-ifp* operon, we designed an optimal sgRNA against the first gene of the operon, *pfp,* using a guide activity predictor^50^. By silencing the first gene in an operon the entire operon is silenced as a polar effect of the CRISPRi mechanism^51^. We performed experiments in both *C. freundii* CF13 and *K. pneumoniae* KPN15. However, RT-qPCR results for *pfp* expression showed that our CRISPRi set-up did not lead to a strong gene silencing in CF13 (Supplementary Fig. 18), so we focused our work on the KPN15 background, where CRISPRi mediated silencing was efficient (Fig. 7B). Our results showed that silencing the *pfp-ifp* operon has no significant fitness effect on the plasmid-free (Tukey’s post-hoc test, coefficient = 4.46, *P* = 0.99) and pOXA-48Δ*lysR*-carrying KPN15 (Tukey’s post-hoc test, coefficient = -30.08, *P* = 0.25; Fig. 7C, Supplementary Fig. 18 and Supplementary Table 11), probably due to the operon’s low level of constitutive expression (Fig. 4D and 7B). In the pOXA-48-carrying strain, on the other hand, silencing the *pfp-ifp* operon produced a clear decrease in fitness (Tukey’s post-hoc test, coefficient = -166., *P* < 0.001; Fig. 7C, Supplementary Fig. 18 and Supplementary Table 11). These results suggest that *pfp-ifp* overexpression increases the fitness of the pOXA-48-carrying strain, which actually shows a higher AUC than the plasmid-free (Tukey’s post-hoc test, coefficient = 84.43, *P* < 0.001; Fig. 7C, Supplementary Fig. 18 and Supplementary Table 11) and pOXA-48Δ*lysR*-carrying strains (Tukey’s post-hoc test, coefficient = -106.61, *P* < 0.001; Fig. 7C, Supplementary Fig. 18 and Supplementary Table 11) in these experimental conditions.

In light of these results, we hypothesized that the overexpression of the *pfp-ifp* operon could be also beneficial for the pOXA-48Δ*lysR*-carrying KPN15 and CF13 strains. To test this hypothesis, we performed growth curves of KPN15 and CF13 carrying both pOXA-48Δ*lysR* and pTet-LysR under different levels of aTc induction (Fig. 7A). Our results showed a clear beneficial effect associated with pTet-LysR carriage in CF13, especially at high induction levels (two-way ANOVA for the interaction between aTc concentration and pTet-LysR, *F*(3,32) = 7.98, *P* < 0.001; for Tukey’s multiple comparisons tests see Supplementary Table 11). For KPN15, we observed an overall growth deficit associated with pTet-LysR (two-way ANOVA for the interaction between aTc concentration and pTet-LysR; *F*(3,32) = 5.43, *P* < 0.004; for Tukey’s multiple comparisons tests see Supplementary Table 11). However, this defect disappeared at the lowest aTc concentration tested (0.075 µg/ml, Tukey’s post-hoc test, coefficient = 45.27, *P* = 0.69), suggesting that this particular level of expression may be beneficial for pOXA-48Δ*lysR*-carrying KPN15.

## Discussion

The complex interactions that occur between plasmids and their hosts shape the evolution of AMR. These interactions can disrupt cellular processes at multiple levels, including the host’s transcriptional regulation. In this work, we explored the impact of the carbapenem-resistance plasmid pOXA-48 on the transcriptomes of 11 MDR clinical enterobacterial hosts. These strains belong to four different clinically relevant species and are natural or sympatric hosts of pOXA-48^9,38^. By integrating transcriptomic analyses with experimental work, we showed that a pOXA-48-encoded transcriptional regulator, LysR_pOXA-48_, mediates a transcriptional crosstalk between the plasmid and the chromosome of *Klebsiella* spp. and *C. freundii* (Figs. 3A and 4A). Specifically, LysR_pOXA-48_ directly increases the expression of a small chromosomal operon comprised of the *pfp* and *ifp* genes (Figs. 3A, 4D and 5). This operon is present in many species of Proteobacteria and shows signs of having been horizontally acquired (Fig. 3B-D). Importantly, our results suggest that the overexpression of the operon leads to a growth benefit for pOXA-48-carrying *K. pneumoniae* KPN15 (Fig. 7C), and could therefore promote plasmid maintenance and transmission in clinical strains.

A growing number of studies have reported plasmid-chromosome crosstalks mediated by plasmid-encoded regulators^10,24^. However, there are only a few documented cases of crosstalks mediated by AMR plasmids^34,36,37^. In this framework, our work is relevant for multiple reasons. First, we studied clinical carbapenemase-producing enterobacteria, which are consistently listed as the top critical group of bacterial pathogens by the World Health Organization^52^. Second, we showed that this crosstalk occurs in multiple MDR enterobacteria of different genera (*Klebsiella* and *Citrobacter*). Third, we performed a thorough analysis of the origin of the *lysR-pfp-ifp* cluster and provided evidence that it is mobilizable, highlighting the relevance of interactions between mobile genetic elements in AMR evolution^12,53^. Finally, we characterized the regulatory activity of LysR_pOXA-48_ and the function of the *pfp-ifp* operon in depth, producing a pOXA-48 *lysR* deletion mutant and combining transcriptomic analyses with CRISPRi silencing. However, our study has limitations. First, due to experimental constraints, we only tested the positive epistatic effect between pOXA-48 and the *pfp-ifp* operon overexpression in one strain. Second, although CRISPRi results suggested that the overexpression of the *pfp-ifp* operon is beneficial in a pOXA-48-carrying strain, we could only partially reproduce the growth benefit associated with the overexpression of the *pfp-ifp* operon in pOXA-48Δ*lysR*-carrying strains. One possible explanation for this result is that the positive interaction between *pfp-ifp* operon overexpression and pOXA-48 only occurs at a narrow range of *lysR*_pOXA-48_ expression. However, it is also possible that there are relevant factors involved in this plasmid-chromosome cross-talk that we are not taking into consideration, such as potential cofactors modulating the function of LysR ^54^. The functional and biological significance of the LysR_pOXA-48_ - *pfp-ifp* operon crosstalk will require further characterization.

Bacterial operons usually encode genes sharing similar or complementary biological functions^55^, and we propose that this could be the case of the *pfp* and *ifp* genes. We showed that the PFP of *K. pneumoniae* KPN15 possesses quercetinase activity (Fig. 6A and Supplementary Figs. 13 and 14). Quercetin is a flavonoid present in many vegetables and is often consumed in the diet^48,56^. Given its iron-chelating properties, quercetin could reduce the bioavailability of iron –a key growth and virulence factor– for gut enterobacteria^56^. We hypothesized that the IFP, being in the same functional family as the EntB isochorismatase, could be involved in the biosynthesis of enterobactin, a high affinity siderophore^49^. Thus, overexpression of the *pfp-ifp* operon by LysR_pOXA-48_ could lead to an increased enterobactin-mediated iron uptake by increasing the biosynthesis of enterobactin (IFP) and the bioavailability of iron –degrading the iron-chelating quercetin (PFP)–. However, we could not demonstrate any growth difference between pOXA-48- and pOXA-48Δ*lysR*-carrying strains in iron-deficient conditions (Supplementary Fig. 16). It is possible that the overexpression of other iron uptake systems observed in pOXA-48Δ*lysR*-carrying strains compared to the pOXA-48-carrying ones is compensating for the presence of the iron chelator (Fig. 6B-C and Supplementary Fig. 15). However, we could not rule out the possibility that the *pfp-ifp* operon has a different function, unrelated to iron uptake. For example, PFP could simply reduce susceptibility to quercetin, which has known antimicrobial properties^48^. Future work will help to uncover the role of this mysterious operon.

In summary, our study uncovers a plasmid-chromosome crosstalk that could facilitate the acquisition of carbapenem resistance in one of the most concerning groups of clinical pathogens, MDR enterobacteria. These results highlight, once again, the key role of plasmids as catalysts for bacterial adaptation beyond being mere drivers of horizontal gene transfer^57^. Given the central role that plasmids play in the dissemination of AMR, characterizing the complex interactions between these mobile genetic elements and their bacterial hosts seems crucial to understanding AMR evolution. Ultimately, we may be able to exploit these interactions to develop new intervention strategies against the increasing threat of AMR.

## Materials and Methods

### Bacterial strains

We selected a subsample of MDR clinical strains from the R-GNOSIS collection, recovered in the Ramón y Cajal University Hospital during an active surveillance-screening programme for detecting and isolating ESBL/carbapenemase-producing Enterobacteriaceae from patients (approved by the Hospital’s Ethics Committee, Reference 251/13)^58^. All strains selected meet the European Centre for Disease Prevention and Control (ECDC) and the Centers for Disease Control and Prevention (CDC)’s conditions for qualifying as MDR Enterobacteriaceae: non-susceptible to ≥1 antimicrobial agent in >3 antimicrobial categories^9,38,59^.

Strains of *Escherichia coli* C325 (ST137), *Citrobacter freundii* CF13 (ST22), *Klebsiella variicola* J57 (ST971) and *Klebsiella pneumoniae* H53 and K147 (ST487 and ST11, respectively) naturally carried plasmid pOXA-48. In a previous work, the genomes of these five strains were sequenced and closed (BioProject PRJNA626430) and isogenic pOXA-48-free clones were obtained by curing the plasmid^39^. Strains of *E. coli* EC10 (ST69) and *K. pneumoniae* KPN04, KPN07 and KPN10 (ST1427) and KPN15 and KPN18 (ST11) did not carry pOXA-48, but were recovered from the same pool of patients, so we defined them as sympatric (ecologically compatible with it). pOXA-48-carrying transconjugants from these strains were obtained before^38,60^. These 11 multidrug resistant strains were representative of the enterobacteria phylogenetic diversity in the R-GNOSIS collection and of pOXA-48 distribution of fitness effects (Supplementary Fig. 1). Additionally, we included the laboratory-adapted *E. coli* MG1655, for which the pOXA-48-transconjugant was obtained before^40^. Experiments and analyses with pOXA-48Δ*lysR* were performed with KPN15 –selected due to belonging to the clinically relevant ST11 and showing a small number of mutations between the pOXA-48-free and transconjugant strain– and CF13. See Supplementary Table 1 for details.

### Plasmids used in this study

The plasmid pOXA-48Δ*lysR* was built by interrupting the pOXA-48-encoded *lysR* gene with an *aph(3’)-Ila* kanamycin-resistance cassette. This plasmid was built using lambda Red recombination as described in^61^. First, a cassette carrying the *aph(3’)-Ila* gene and 500 bp left and right homology regions was cloned into plasmid pMP7^62^ using Gibson Assembly. The cassette was amplified from the resulting pMP7-HR-kan-HR plasmid. The primers used in this construction and a more complete description on how pMP7-HR-kan-HR was built can be found in Supplementary Table 1.

The pTet-LysR plasmid was built using Gibson Assembly. The primers can be found in Supplementary Table 1.

The plasmid pFR56apm was used for expression of the CRISPRi machinery. This plasmid is based on pFR56^63^, with the chloramphenicol-resistance marker substituted by an apramycin-resistance gene. pFR56apm expresses dCas9 under the inducible promoter pHlf, and the single guide RNA + scaffold under a constitutive promoter. Two versions of the plasmid were used in this study: (i) a non-targeting control, with 20 random nucleotides as sgRNA that don’t target anything in the chromosome of the strains, and (ii) a version that allows for silencing of the *pfp-ifp* operon, constructed as described in the original pFR56 paper and chosen out of all possible guides against *pfp* using a guide activity predictor^50^. The primers used for the construction of the pFR56apm versions are described in Supplementary Table 1.

### Genomic analyses

#### Whole genome sequencing

The genomes of the six pOXA-48 clinical transconjugants and the *E. coli* MG1655 transconjugant were sequenced with long-read technology to generate closed sequences to use as references for the transcriptomic analyses. DNA extraction was performed as in^64^. Genomic DNA was sequenced at the Microbial Genome Sequencing Center (MiGS; Pittsburgh, PA, USA) using Nanopore technology or in-house with a MinION Mk1B as in^64^. Illumina reads of the six clinical transconjugants were obtained before (BioProject PRJNA641166)^38^ or were resequenced at MiGS with NextSeq 2000, generating 150 bp paired-end reads. The pOXA-48-carrying MG1655 strain was sequenced at MicrobesNG (Birmingham, UK) with a MiSeq, generating 250 bp paired-end reads.

#### Closing reference genomes

Quality control of Illumina reads was performed with FastQC v0.11.9 (https://www.bioinformatics.babraham.ac.uk/projects/fastqc/). Illumina reads of the six clinical transconjugants were processed with Trim Galore v0.6.4 (Cutadapt v2.8, https://github.com/FelixKrueger/TrimGalore) to trim ends with Phred quality scores lower than 20 and remove adapters and reads shorter than 50 bp. Reads of pOXA-48-carrying MG1655 were pre-processed by MicrobesNG. Hybrid assemblies were obtained with Unicycler v0.4.9^65^ using the Illumina reads from^38^ and MicrobesNG and the Nanopore reads from MiGS. The assemblies of pOXA-48-carrying KPN04, KPN07 and KPN18 were fragmented, possibly due to short- and long-reads originating from different DNA extractions. Thus, Illumina reads from MiGS and long-reads generated in-house were used to generate new hybrid assemblies with Unicycler. These assemblies were complete except for a plasmid that could not be closed in pOXA-48-carrying KPN04 and KPN18. To close the plasmid, Nanopore reads were processed with filtlong v0.2.1 (https://github.com/rrwick/Filtlong) to obtain a subset of high identity reads with a minimum read depth of 85x and filtered reads were assembled with Flye v2.9^66^. Resulting assemblies were polished with Medaka v1.4.3 (https://github.com/nanoporetech/medaka) and several rounds of Pilon v1.24^67^, mapping the trimmed Illumina reads until no more changes were made. Finally, polished assemblies were recircularized with circlator v1.5.5^68^. This way, the incomplete plasmids from the Unicycler assemblies were replaced by the closed polished plasmids in the FASTA files. In pOXA-48-carrying EC10 and MG1655, the assembly of pOXA-48 was missing a 760 bp region affecting the first copy of IS1 and a 90 bp region affecting the *traC* gene, respectively. However, these represent assembly errors because there were reads mapping to these regions when using the pOXA-48_K8 variant (accession number MT441554) as reference. Final reference genomes were annotated with PGAP v2021-07-01.build5508^69^.

#### Phylogenetic tree of enterobacterial strains

Mashtree v1.2.0^70^ was used to generate a phylogenetic tree of mash distances between the closed genomes of the 12 pOXA-48-carrying strains, using a bootstrap of 100. The tree was represented with the Interactive Tree of Life (iTOL)^71^ online tool.

### Transcriptomic analyses

#### RNA extraction, sequencing and quality control

RNA extraction was performed as in^64^. Briefly, cells were grown with continuous shaking in LB medium. When cultures reached an OD_600_ of 0.5 (exponential phase), cells were collected at 4°C, pelleted by centrifugation at 4°C 12,000 g for 1 min and frozen immediately at -70°C. Total RNA was purified from each sample using the NZY Total RNA Isolation kit (NZYTECH). RNA concentration was determined using the Qubit RNA Broad-Range Assay following the manufacturer’s instructions. Additionally, the quality of the RNA was examined using the Tape-Station system (Agilent). RNA was ribodepleted and sequenced at the Wellcome Trust Centre for Human Genetics (WTCHG; Oxford, UK) using the Illumina’s NovaSeq6000 platform, resulting in >8 million reads per sample (first RNA-Seq dataset). Three biological replicates per strain and condition (carrying and not carrying pOXA-48) were sequenced, except for strains CF13, KPN10 + pOXA-48, KPN10 and MG1655 that only two replicates were sequenced due insufficient RNA Integrity number (RIN score). To further confirm and characterize the plasmid-chromosome crosstalk, another RNA extraction and sequencing was performed for strains CF13 and KPN15 carrying pOXA-48 or the pOXA-48Δ*lysR* variant (see the RT-qPCR Methods for the RNA extraction protocol). Total RNA (three biological replicates for each strain and condition) was ribodepleted and sequenced with the NextSeq2000 platform at SeqCoast Genomics (Portsmouth, NH, USA), generating >6 million reads per sample (second RNA-Seq dataset). The quality of the reads was checked with FastQC v.0.11.9. Reads were trimmed, adapter-removed and filtered for length lower than 50 bp using Trim Galore v0.6.4.

#### Differential expression analysis

Trimmed reads were mapped to their corresponding reference genome with BWA-MEM v0.7.17^72^. The percentage of mapped reads was >99% for all samples. The appropriate presence or absence of pOXA-48 in the replicates was confirmed by inspecting the read alignments. A replicate in pOXA-48-cured K147 and CF13 (first RNA-Seq dataset) actually carried the plasmid, so they were included as an additional replicate of the pOXA-48-carrying strains. The function *featureCounts* from the Rsubread v2.14.2 package^73^ was used to obtain raw counts of reads mapping to each genomic feature (including CDS, ncRNA, tRNA, tmRNA, antisense RNA, RNase P and SRP RNA), with strand specificity. The percentage of alignments successfully assigned to features was >85%. The presence of all resident plasmids was confirmed by reads mapping to their respective references (raw counts). DE analysis was performed using DESeq2 v1.40.1^74^ from raw counts by comparing the pOXA-48-carrying strains against their respective pOXA-48-free strains (first RNA-Seq dataset) or pOXA-48Δ*lysR*-carrying strains (second RNA-Seq dataset). Raw DEGs (log_2_FC cutoff of 0 and adjusted *p* value < 0.01) were filtered to obtain significant DEGs with an adjusted *p* value < 0.05. Raw counts and significant DE results are available at S4 and Supplementary Table 10s. For the interpretation of the DE of pOXA-48 genes in the comparison pOXA-48 *vs*. pOXA-48Δ*lysR* (second RNA-Seq dataset), genes nearing the P*_cat_*-*aph(3’)-IIa* insertion, including the *lysR* gene, were not considered due to possible polar effects. The Operon-mapper webserver (January 2023)^75^ was used to determine whether the *pfp* and *ifp* genes formed a single operon.

#### Transcripts per million (TPM) calculation

TPM values were calculated from FPKM values computed with DESeq2 from raw counts. To plot pOXA-48 gene expression, TPM values of pOXA-48 genes were normalized by the median chromosomal TPM, since it shows lower coefficient of variance across samples (CV = 0.22) than the housekeeping genes *rpoD* (CV = 0.39), *dnaE* (CV = 0.50) or *recA* (CV = 0.46). TPM values are available at Supplementary Table 3.

#### Gene set enrichment analysis (GSEA)

Raw DEGs from the first RNA-Seq dataset were annotated with Gene Ontology (GO) terms for biological processes (BP), molecular functions (MF) and cellular components (CC), retrieved from the UniProt database^76^ on May 2023 by mapping the RefSeq IDs from the PGAP annotation files to UniProtKB IDs. The percentage of raw DEGs that were annotated with GO terms was >63%. GSEA was performed for each strain on the lists of chromosomal and plasmid raw DEGs, separately, ranked by log_2_FC (shrunk with DESeq2’s *apeglm* method) with clusterProfiler v4.8.1^77^, correcting for multiple tests with the Benjamini-Hochberg procedure (Supplementary Table 5).

#### Variant calling

To assess whether mutations accumulated in the chromosome or other plasmids during growth or construction of pOXA-48-, pOXA-48Δ*lysR*-carrying or pOXA-48-free samples affected gene expression, variant calling was performed on the transcribed regions with Snippy v4.6.0 (https://github.com/tseemann/snippy), using the RNA-Seq mappings obtained previously (--bam). Only mutations not present in all replicates per strain were considered, since mutations in all samples of a strain would not affect differential expression (Supplementary Table 2). Of the total 80 SNPs identified in transcribed regions, only 12 showed DE of the mutated gene (Supplementary Table 2). To confirm or discard suspected SNPs in plasmid pOXA-48, the RNA-Seq reads were mapped to the sequence of the pOXA-48_K8 variant (accession number MT441554), the variant that was introduced to the transconjugants (Supplementary Table 2).

#### Clustering of strains

To further analyze common transcriptomic responses between samples, 2,488 single copy orthologs (SCO; orthogroups which contain only one gene per strain) were identified with OrthoFinder v2.5.4^78^. PCA (Supplementary Fig. 3A) and t-SNE (R package *Rtsne*, perplexity = 5; Supplementary Fig. 3B) were performed on the SCO of pOXA-48-carrying and -free replicates to visualize patterns of response to carriage of the plasmid. For this, total raw counts were first normalized with VST (variance stabilizing transformation; DESeq2 package) and subsetting of the SCO was performed afterwards. Hierarchical clustering (complete linkage method) was performed to find strains with similar DE profiles (Supplementary Fig. 3C).

### Analysis of the origin of the *lysR-pfp-ifp* cluster

#### *Identification of the* lysR-pfp-ifp *cluster in a database of Proteobacteria*

To explore the origin of the *lysR-pfp-ifp* cluster, the protein sequences of the LysR, PFP and IFP were first searched in STRING^79^. The conserved neighborhood view revealed the *lysR-pfp-ifp* cluster is present in different species across the Proteobacteria phylogeny (Supplementary Fig. 7). To further analyze the origin of the *lysR-pfp-ifp* cluster, all Proteobacteria RefSeq genomes with a “chromosome” or “complete” assembly level were downloaded from the NCBI database (retrieved on July 14th, 2023 from https://www.ncbi.nlm.nih.gov/datasets/genome/), which comprises 16,219 Gammaproteobacteria, 2,878 Betaproteobacteria, 2,441 Alphaproteobacteria, 1,086 Epsilonproteobacteria, 5 Deltaproteobacteria and 2 Zetaproteobacteria (Supplementary Table 7). A MacSyFinder v2.0 model^80^ was constructed to identify the *lysR-pfp-ifp* cluster in the Proteobacteria database. To construct the model, the Pfam HMM profiles (release 35.0)^81^ of the three proteins (LysR: PF03466, PFP: PF02678, IFP: PF00857) were downloaded from InterPro v95.0 after an InterProScan 5 search^82^. The model, available at https://github.com/LaboraTORIbio/RNA-Seq_enterobacteria_pOXA-48, is configured to find systems that contain the three genes consecutively (min_mandatory_genes_required = 3, min_genes_required = 3, inter_gene_max_space = 0), although order and sense cannot be specified. Then, the model was used with MacSyFinder v2.1.1 to find the *lysR-pfp-ifp* cluster in the database. The MacSyFinder results were filtered using in-house scripts to remove systems with incorrect gene order and orientation. Correct systems were defined as having the gene order *lysR-pfp-ifp* or *lysR-ifp-pfp,* with *pfp* and *ifp* in the same strand and *lysR* in antisense, so that *lysR* is likely to control the expression of *pfp* and *ifp*. Also, in strains with more than one system detected (24 Gamma-, 32 Beta- and 18 Alphaproteobacteria), only the system encoding the *lysR* with higher percentage of protein sequence identity with the *lysR* of pOXA-48 (BLASTp v2.11.0 search^83^) was kept. After filtering, 5,983 Gamma-, 191 Beta- and 172 Alphaproteobacteria strains encoded the *lysR-pfp-ifp* cluster.

#### *Phylogenetic tree of the* lysR-pfp-ifp *cluster*

To reduce the size of the database of Proteobacteria encoding the *lysR-pfp-ifp* cluster, the concatenated protein sequences of the cluster (in the order *lysR-pfp-ifp*) were clustered using USEARCH v11.0.667^84^ with a 0.99 identity threshold to avoid losing evolutionary information. The LysR, PFP and IFP protein sequences of the 698 representative strains of each resulting cluster were aligned with MAFFT v7.453^85^. Alignments were trimmed with trimAl v1.4^86^ and concatenated with catfasta2phyml (https://github.com/nylander/catfasta2phyml) for constructing the tree of the *lysR-pfp-ifp* cluster. Four phylogenetic trees (*lysR-pfp-ifp* cluster [Fig. 3B], LysR [Supplementary Fig. 8], PFP [Supplementary Fig. 9] and IFP [Supplementary Fig. 10]) were constructed with IQ-TREE v1.6.12^87^ with best evolutionary model selection and 1,000 ultrafast bootstrap. Trees were visualized and edited with iTOL^71^.

#### Ancestral character reconstruction (ACR)

To construct a phylogenetic tree of Gamma-, Beta- and Alphaproteobacteria for ACR, including strains encoding and lacking the *lysR-pfp-ifp* cluster, the database had to be reduced. The 698 representative *lysR-pfp-ifp* cluster-encoding strains (comprising 321 different species) were selected. From these 321 species, 138 species included strains encoding or lacking the *lysR-pfp-ifp* cluster. To include these events in ACR, 102 random strains from 102 species that had <10 strains not encoding the *lysR-pfp-ifp* cluster, and 180 random strains from 36 species (5 per species) that had ≥10 strains not encoding the *lysR-pfp-ifp* cluster were selected. Additionally, 35% of the remaining strains lacking the *lysR-pfp-ifp* cluster were selected (359 Gamma-, 131 Beta- and 234 Alphaproteobacteria). Finally, 10 random Epsilonproteobacteria strains were selected as outgroups. Thus, in total, 1,714 strains were selected. The phylogenetic tree of Proteobacteria was constructed from 128 conserved bacterial single-copy genes, described in^88^ and curated in^89^. These genes were identified in the 1,714 genomes using the Pfam HMM profiles (release 35.0) with a HMMER v3.3 (option --cut_ga)^90^ search. Hits were filtered by score using the cutoffs reported in^88^, resulting in 125 gene markers present in >85% of strains. Protein sequences of each family were aligned with MAFFT (option --auto) and alignments were trimmed and concatenated with trimAl and catfasta2phyml. The tree was constructed with IQ-TREE as before (-m MFP -bb 1000) and rooted at the Epsilonproteobacteria clade with iTOL. Four rogue taxa were identified with TreeShrink v1.3.9^91^ and iTOL was used to prune the tree, removing the rogue taxa and 14 Gammaproteobacteria strains (none encoding the *lysR-pfp-ifp* cluster) that branched outside of the Gamma/Betaproteobacteria clade (Supplementary Fig. 19). ACR was performed with PastML v1.9.34^92^ with default parameters, using the pruned tree and a table of presence/absence of the *lysR-pfp-ifp* cluster (Supplementary Fig. 11).

#### *Genomic neighborhood of the* lysR-pfp-ifp *cluster*

To analyze the regions upstream and downstream the *lysR-pfp-ifp* cluster, a subset of relevant and representative species was selected: 19 Enterobacteriaceae strains (including the Enterobacteriaceae strains that branched outside the Enterobacteriaceae clade, Fig. 3B) and other 5 Gamma-, 4 Beta- and 3 Alphaproteobacteria strains (Supplementary Fig. 12). Except for the rogue Enterobacteriaceae strains and the strains used in this study, the remaining strains were randomly selected. Strains from the same species were selected so that they were distant on the phylogenetic tree (Fig. 3B). First, the regions 10 Kbp upstream and 10 Kbp downstream of the *lysR-pfp-ifp* cluster were extracted using the samtools v.1.14 *faidx* command^93^. To obtain annotation files compatible with clinker, the fasta files of these regions were annotated with Prokka v.1.14.6^94^ using the original annotated proteins as first priority (--proteins flag). The GBK files provided by Prokka were used as input for clinker v.0.0.28^95^, executed with default parameters (i.e. minimum alignment sequence identity = 0.3). Two static HTML documents were plotted: one for all the 31 strains selected (Supplementary Fig. 12), and another with non-redundant representative strains (n = 8; Fig. 3D). In Fig. 3D, genes were colored based on the main functional groups identified (e.g. iron uptake) as well as on the potential relatedness to HGT (tRNAs and transposases).

#### CG content and codon adaptation index (CAI)

The GC content and CAI was computed for the subset of non-redundant *lysR-pfp-ifp* cluster-encoding Proteobacteria (n = 8) to assess whether the *lysR-pfp-ifp* cluster showed further signs of horizontal acquisition. The GC content of the chromosome and the *lysR-pfp-ifp* cluster (from the start of *lysR* to the end of the second operon gene) were calculated as the ratio of GC nucleotides by the total nucleotides of the region (Supplementary Table 8). Codon usage compatibility was assessed by comparing the CAI of the *lysR-pfp-ifp* cluster genes to the median CAI of genes encoding for ribosomal proteins, which are highly expressed and thus have an optimized codon usage. First, a codon usage table was created from genes encoding for ribosomal proteins (retrieved from annotation files searching for the term “ribosomal protein”) using the *cusp* function of EMBOSS v6.6.0.0^96^. The CAI of each *lysR-pfp-ifp* cluster gene and each ribosomal protein gene was computed using the *cai* function of the EMBOSS package, using the codon usage table previously created (Supplementary Table 8).

### RT-qPCR

#### Bacterial RNA extraction

Overnight bacterial cultures grown in LB were diluted to 1:1000 in fresh medium. When necessary, LB medium was supplemented with anhydrotetracycline (aTc) 0.3 µg/mL or 2,4-diacetylphloroglucinol (DAPG) 50 μM. Cultures were grown in aerobic conditions at 37 °C with agitation until an OD_600_ of 0.2-0.3 was reached. Then, 3 mL of each culture were pelleted (11,000 rpm, 1 min) to perform total RNA extraction (using BioRad’s Aurum™ Total RNA Mini Kit). RNA concentration and purity were assessed using a nanodrop and all samples shown to have A260/A280 and A260/A230 ratios ≥ 2. RNA integrity and lack of contamination by residual genomic DNA were evaluated by electrophoresis in a 1% agarose gel. All samples had intact 16S and 23S ribosomal RNA. Two or three biological replicates were obtained per group.

#### Reverse transcription (RT)

RT was carried out for each replicate using 1 μg of RNA as template and the High-Capacity cDNA Reverse Transcription Kit with RNase Inhibitor from Applied Biosystems.

#### qPCR

To quantify expression levels of *lysR*_pOXA-48_, *bla*_OXA-48_, *pfp* and *rpoB* (endogenous control), 2 μL of the previous undiluted cDNA were used as template in a 20 μL-reaction containing NZYSupreme qPCR Green Master Mix from Nzytech and the specific primer-pairs for each of the genes (Supplementary Table 1). Fluorescence data was measured using a 7500 Real-Time PCR System from Applied Biosystems. Relative mRNA quantities were calculated using the comparative threshold cycle (Ct) method and normalized using *rpoB* gene expression (2^-ΔCt^). Efficiencies of each reaction were calculated by the standard curve method in triplicate (*bla*_OXA-48_: Efficiency = 0.95, R2 = 0.99; *lysR*_pOXA-48_: Efficiency = 0.98, R2 = 0.99; *pfp*: Efficiency = 0.97, R2 = 0.99; *rpoB*: Efficiency = 0.98, R2 = 0.99). Two independent experiments in triplicate were performed.

### Evaluation of the antimicrobial effect of quercetin by the broth microdilution method

The susceptibility to quercetin of pOXA-48-carrying strains versus pOXA-48Δ*lysR*-carrying strains was evaluated using the broth microdilution method. First, a quercetin dihydrate 10 mg/mL stock solution was prepared (97% purity Quercetin dihydrate from Fisher Scientific dissolved in DMSO from Sigma). Then, two-fold serial dilutions were performed in MH medium, starting at 128 µg/mL and down to 1 µg/mL. 100 µL of the different dilutions were added per well to a 96-well microtiter plate. On top of that, a further 100 µL of bacterial inoculum was added per well, obtained from a 1:2000 dilution of an overnight culture in MH medium (5 x 10_5_ CFU/mL). The following controls were included: (i) a sterility control per dilution, where no bacterial inoculum was added, and (ii) a DMSO control prepared in the same way as the quercetin dilutions, to assess for the toxicity of DMSO alone. Growth curves (1 replicate for DMSO and 2 replicates for DMSO+QC, per concentration and strain) were performed by growing the cultures in a plate reader BioTek SynergyHTX for 24 h, at 37 °C with orbital shaking (282 cpm) and measuring the OD_600_ every 10 min.

### Iron chelator assays

Growth curves of plasmid-free, pOXA-48-carrying and pOXA-48Δ*lysR*-carrying KPN15 and CF13 were performed in the presence of the Iron Chelator 2,2-Bipyridyl from VWR (ref. 30569.09). Concentrations of 2,2-Bipyridyl ranging from 0.016 mM to 0.5 mM were tested. Curves were performed from overnight cultures grown without 2,2-Bipyridyl and diluted down to 1:1000. Growth curves (3 replicates per condition and strain) were performed for 24 h, using a plate reader BioTek SynergyHTX, at 37 °C with orbital shaking (282 cpm) and measuring the OD_600_ every 10 min.

### Growth curves expressing LysR_pOXA-48_ in *trans*

Growth curves were performed to assess the effect of the overexpression of LysR_pOXA-48_ in plasmid-free KPN15 and CF13, as well as their pOXA-48- and pOXA-48Δ*lysR*-carrying counterparts. Overexpression was performed in *trans* using pTet-LysR (LysR_pOXA-48_ under the control of the pTet promoter). Growth curves were started by diluting overnight cultures of each strain to a factor of 1:1000, in LB and LB supplemented with a range of aTc concentrations (0.075, 0.15 and 0.3 µg/mL). Cultures were grown in a BioTek SynergyHTX plate reader, at 37°C, for 24h, measuring OD_600_ every 10 min. Measurements were performed for 5 replicates of each strain/induced pairing.

### Gene silencing through CRISPR interference (CRISPRi)

The two versions of pFR56apm (non-targeting control and the one targeting the *pfp-ifp* operon) were transformed into *E. coli* β3914ø + pTA-MOB^97,98^. This strain is auxotrophic for diaminopimelic acid (DAP) and can be used as a counter-selectable donor strain. pTA-MOB allows for mobilization of pFR56, which carries an ori*T* but lacks the genes necessary for conjugation. This set-up was used for conjugation of the two versions of pFR56apm into plasmid-free CF13 and KPN15, and their pOXA-48- and pOXA-48Δ*lysR*-carrying counterparts.

The resulting strains were grown overnight in LB supplemented with apramycin (50µg/mL), to select for pFR56, and DAPG (50µM), to induce *dcas9* expression. In the morning, an 1:1000 dilution was performed for each strain in LB supplemented with apramycin and DAPG, and cultures were grown in a BioTek’s SynergyHTX plate reader, at 37°C, for 24h, measuring OD_600_ every 10 min. Measurements were performed for 16 replicates of each strain.

### Statistical analyses

All statistical analyses were performed with R v4.3.1 or v4.4.0 (www.R-project.org). Packages *ggplot2*, *ggpubr*, *ggfortify*, *cowplot*, *scales*, *pheatmap*, *enrichplot*, *RColorBrewer*, *reshape2* and *tidyverse* were used for data representation and manipulation. R base and *car* packages were used for statistical tests. Normality and homoscedasticity of the data was assessed with the Shapiro–Wilk and Levene tests, respectively. Spearman’s rank correlation was used for non-normal data. To compare means in the RT-qPCR results (see Fig. 4D), a one-way ANOVA (for KPN15, *F*(2,3) = 165.4, *P* = 0.0009; for CF13, *F*(2,3) = 82.4, *P* = 0.002) with Dunnett’s post hoc test, or *t*-tests were used when appropriate. The area under the curve (AUC) of growth curves was computed with the R package *flux* v0.3-0.1. To compare growth differences in the pTet-LysR and CRISPRi assays (Fig. 7), two-way ANOVA and the post hoc Tukey’s tests were performed (Supplementary Table 11). To assess the effect of the concentration of the drugs DMSO or DMSO+QC in growth (AUC, as the dependent variable), two GAMs were constructed for each strain (due to observable differences in growth and drug response) with the R package *mgcv* v1.9-1. A non-linear relationship was fitted per genotype (pOXA-48 or pOXA-48Δ*lysR*) and drug (DMSO or DMSO+QC) interaction, including the interaction terms for both explanatory variables (formula: *AUC ∼ s(concentration, k=5, by=genotype_drug) + genotype * drug*). Genotype pOXA-48Δ*lysR* and drug DMSO+QC were set as reference levels to obtain relevant comparisons. The Restricted Maximum Likelihood (REML) method was used. The optimal number of basis functions (*k*) was selected by comparing the model fit statistics (adjusted R-square, deviance explained and -REML; Supplementary Table 11).

## Data and code availability

Closed reference genomes are available at BioProjects PRJNA626430 and PRJNA1071971 (see Supplementary Table 1). Genomic and transcriptomic sequencing data generated in this study is available at BioProject PRJNA1071971. Raw data generated during DE analysis can be found at the GEO Series GSE269852. Detailed bioinformatics workflows and the code produced in this study are available at https://github.com/LaboraTORIbio/RNA-Seq_enterobacteria_pOXA-48. Experimental raw data can be found in Source Data.

## Acknowledgements

We thank David Bikard and the Institut Pasteur for the gift of plasmid pFR56.

## Author contributions

L.T.C., A.C.V., C.H., A.F.C. and A.S.M. conceptualized the study. L.T.C., A.C.V., C.H., A.F.C. and A.S.M. designed the methodology. L.T.C. and J.S.D. performed the bioinformatic analyses. A.C.V, C.H., A.A.dV., S.Q. and A.F.C. performed the experiments. L.T.C., A.C.V., J.S.D., A.F.C. and A.S.M. contributed to data analysis. D.M. supervised experimental work. E.R. supervised bioinformatic analyses. A.S.M. was responsible for funding acquisition and supervision. L.T.C., A.C.V., A.F.C. and A.S.M. wrote the original draft and undertook the reviewing and editing process. All authors supervised and approved the final version of the manuscript.

## Supplementary Information for the manuscript

### Supplementary Results and Discussion

#### Expression of other resident plasmid’s genes

We analyzed the differential expression (DE) of genes of other resident plasmids (excluding pOXA-48) in the 11 clinical strains. To find groups of genes with similar functions that tend to be DE in the same direction, we performed gene set enrichment analysis (GSEA) on the sets of plasmid genes. Transposases or insertion sequences (ISs) were enriched in three *Klebsiella* spp. strains (Supplementary Table 5). In J57, transposition processes tended to be upregulated with pOXA-48 carriage, while in KPN07 and KPN10, downregulated. The IS families involved in these patterns were diverse in the three strains (Supplementary Table 5). Interestingly, we previously observed a similar pattern with the pOXA-48-encoded IS1, where IS1 was downregulated in strains encoding multiple copies of IS1, like in KPN07 and KPN10 due to high PCN, and upregulated in strains encoding few copies of IS1, like J57, which does not encode IS1 in the chromosome or other plasmids^1^. This suggests that the IS1 of pOXA-48 could be influencing the expression of other IS families. Additionally, processes involved in response to antibiotics were upregulated in KPN15 (Supplementary Table 5).

#### Analysis of the conservation of LysR_pOXA-48_

If overexpressing the operon is important for pOXA-48-carrying strains, LysR_pOXA-48_ could be under selective pressure to maintain its regulatory function, and thus its protein sequence should be conserved. To study this, we analyzed the protein sequences of LysR_pOXA-48_ in the RefSeq database of Proteobacteria, using the LysR_pOXA-48_ sequence of pOXA-48_K8 –the most frequent variant isolated in our hospital– as query. We filtered BLASTp searches by 90% identity and full alignment length (303 amino acids) to discard matches with other LysR regulators (Supplementary Table 12). This way, we identified LysR_pOXA-48_ only in Gammaproteobacteria (288 strains). Importantly, there were no hits in the range 90-96% of identity, indicating high sequence conservation. Furthermore, when lowering the alignment length threshold to >250 amino acids, only five additional hits were obtained, supporting the high sequence conservation.

We used ABRicate v1.0.1 (https://github.com/tseemann/abricate) with the PlasmidFinder database to determine whether LysRs with high identity with LysR_pOXA-48_ were encoded on plasmids or chromosomes. We performed the analyses separating hits in two groups: (i) LysRs with 100% identity with LysR_pOXA-48_, and (ii) LysRs with <100% (and >96%) identity with LysR_pOXA-48_. Most LysR_pOXA-48_ (71%) were encoded on plasmids, specially on pOXA-48-like plasmids (n = 172). Chromosomally-encoded LysR_pOXA-48_ were mainly present in *Shewanella* spp., from where the *bla*_OXA-48_-*lysR* genes were originally transposed to the IncL plasmid backbone. Most plasmid-encoded LysR_pOXA-48_ (93%) shared 100% with the reference sequence, and 90.6% were encoded in pOXA-48-like plasmids (Supplementary Table 12). Then, only 13 LysR_pOXA-48_ shared less than 100% identity with the reference, and 38% of these were encoded in other plasmid backbones (Supplementary Table 12).

To analyze amino acid substitutions in LysRs encoded on plasmids, we aligned the protein sequences of these LysRs with MAFFT v7.453. Alignments were visualized in ESPript v3.0^2^. We used the SIFT^3^ and Mutpred2^4^ webservers (May 2024) to assess the impact of amino acid substitutions on protein function. The 13 LysR_pOXA-48_ with less than 100% identity with the reference shared the same amino acid substitution (S112G) compared to the reference LysR_pOXA-48_ (Supplementary Fig. 20), and it was predicted to not affect protein function (SIFT score = 0.45, Mutpred2 score = 0.36), supporting the protein sequence of LysR_pOXA-48_ is under high selective pressure.

Next, we calculated the ratio of non-synonymous to synonymous nucleotide substitutions (N/S) of LysR_pOXA-48_ among the 172 pOXA-48-like plasmids. For this, we used the nucleotide sequence of pOXA-48_K8 (accession number MT441554), the most common pOXA-48 variant found in clinical isolates from the Hospital Universitario Ramón y Cajal^5^, as a common reference to analyze the variants. We employed the NCBI PGAP pipeline v.2022-12-13.build6494 to annotate the reference sequence. We used Snippy v.4.6.0 (https://github.com/tseemann/snippy) to identify the variants from each of the plasmid sequences downloaded from RefSeq using the contigs as input. We merged the mutations by nucleotide position for each of the genes (i.e. different n plasmids with the same SNP were identified as 1 event). We then classified the identified variants by their effect as Non-synonymous ("missense_variant") or Synonymous ("synonymous"). Finally, we calculated the ratio of Non-synonymous over Synonymous SNPs (N/S) for each of the plasmid genes as a proxy for the strength of selection. We identified a total of 599 different variants (Supplementary Table 12). Only one of these variants affected *lysR*_pOXA-48_ (Non-synonymous: Ser112Gly), which correspond to the eight pOXA-48-like plasmids encoding for the LysR_pOXA-48_ with the S112G amino acid substitution (Supplementary Table 12). Given the low number of mutations in *lysR*_pOXA-48_, the N/S (1/0) could be misleading (Supplementary Table 12).

Thus, we further tested whether mutations were equally distributed along the plasmid genes by performing a permutation test (n permutations per gene = 10,000). First, we corrected the number of mutations per gene by the gene length. Then, we used as test statistic the absolute difference between the number of mutations per base for each gene (Ng) and the mean Ng for the rest of the genes of the plasmid (Ng-Np). In each permutation, an Ng was randomly sampled and compared with the mean of the rest of the genes. The *p*-value would be the probability of a random value being more extreme than the observed value for each gene. We found four genes significantly enriched (*P* < 0.05) in the number of mutations per base (*traR*, a helix-turn-helix transcriptional regulator, and two hypothetical proteins, Supplementary Table 12), but we did not detect any significant differences for *lysR*_pOXA-48_ (*P* = 0.13, Supplementary Table 12). However, when analyzing the number of mutations per base for all pOXA-48 genes, *lysR*_pOXA-48_ was the second least targeted gene, after *traY* (Supplementary Table 12).

These results, together with the fact that *lysR*_pOXA-48_ was the only gene transposed along with *bla*_OXA-48_ from the chromosome of *Shewanella* spp. to pOXA-48, suggest that the function of LysR_pOXA-48_ is important for pOXA-48 carriers and thus is under selective pressure to retain its regulatory function, possibly over the *pfp-ifp* operon genes.

## Supplementary Figures

**Supplementary Fig. 1.**
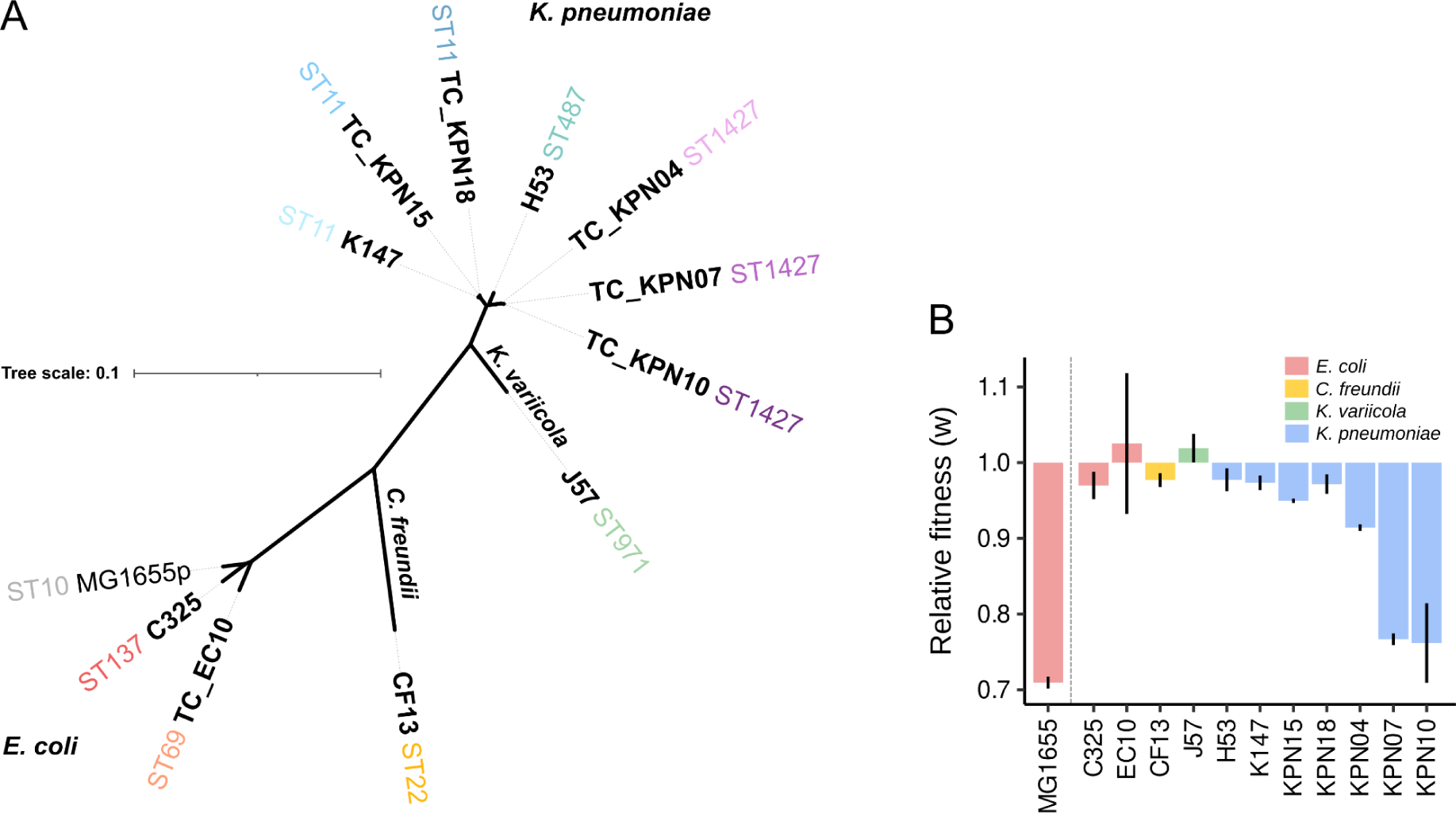
Enterobacterial strains. (A) Unrooted phylogeny of mash distances between the 11 clinical enterobacterial strains, marked in bold, and the laboratory *E. coli* strain MG1655. Trees were constructed with the closed genomic sequences of the pOXA-48-carrying strains. Sequence Types (STs) are indicated. (B) Distribution of fitness effects of pOXA-48-carrying strains. Relative fitness (*w*) values were previously obtained from competition experiments between pOXA-48-carrying and their respective pOXA-48-free strains^38–40^. Values >1 indicate that pOXA-48 carriage is associated with a fitness advantage, while values <1 indicate a reduction in fitness associated with pOXA-48. Error bars indicate the standard error of the mean.

**Supplementary Fig. 2.**
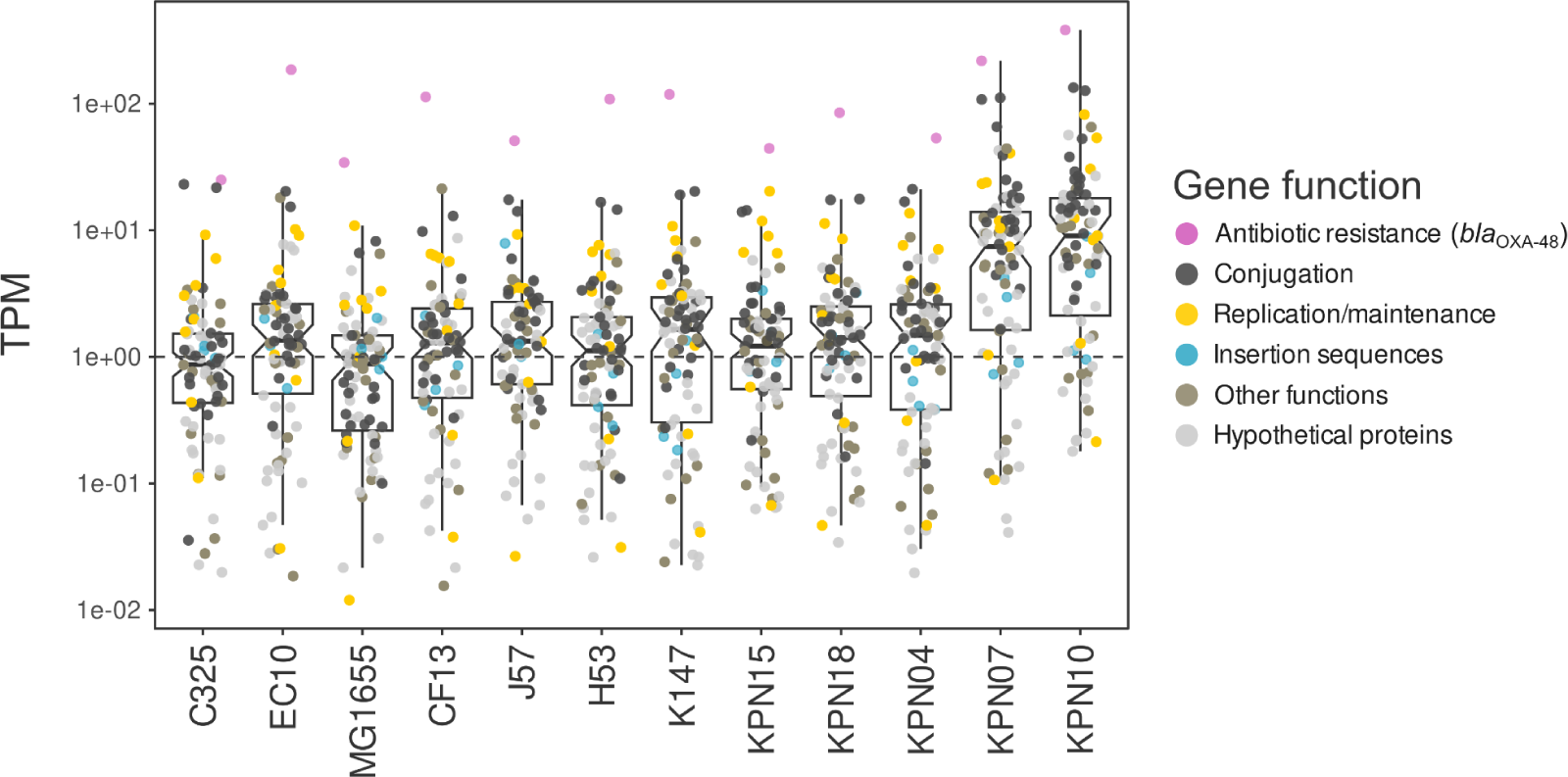
Expression of pOXA-48 genes. Boxplots of the expression of pOXA-48 genes of each strain, measured as the median TPM (Transcripts Per Million) between replicates, normalized by median chromosomal TPM values of the corresponding strain (horizontal dashed line). The *y* scale is log_10_-transformed for visualization. TPM > 1 indicates that pOXA-48 genes are expressed more than the median expression of chromosomal genes, while TPM < 1 indicates that pOXA-48 gene expression is below the median chromosomal TPM values. Median TPM values are indicated as horizontal lines inside the boxes and the notches represent the 95% confidence interval for the median. The upper and lower box hinges correspond to the 25th and 75th percentiles and the whiskers extend to observations within 1.5x the interquartile range.

**Supplementary Fig. 3.**
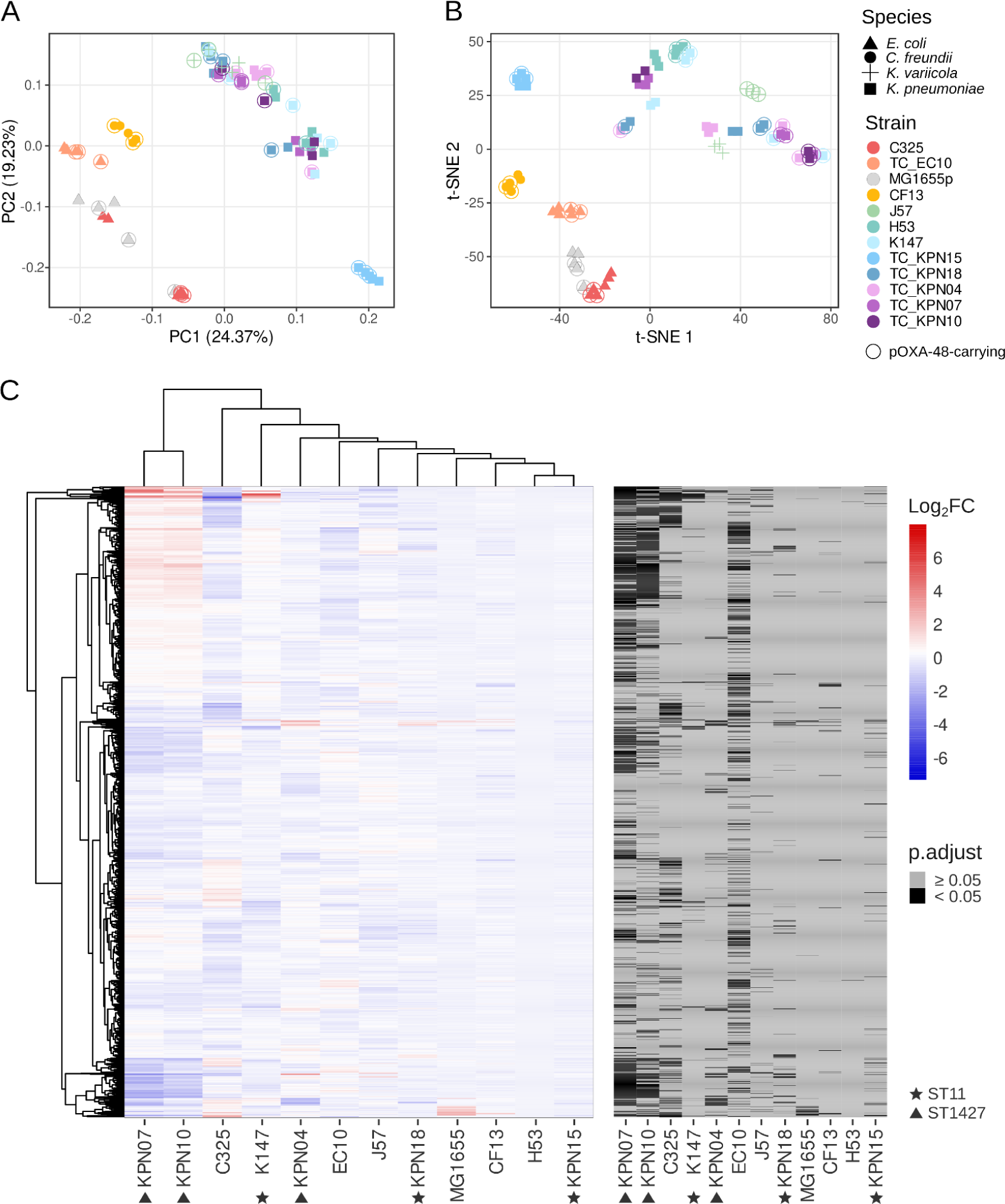
Strain-specific expression of 2,488 single copy orthologs (SCO). (A) Principal component analysis and (B) t-distributed stochastic neighbor embedding of pOXA-48-free and -carrying (indicated with circles) enterobacterial replicates. Both panels represent gene expression of 2,488 single copy orthologs (see Methods). Shapes and colors indicate the species and the strain of each sample, respectively. (C) The heatmap on the left panel represents the differential expression (log_2_FC) of 2,488 SCO calculated by comparing pOXA-48-carrying strains to their pOXA-48-free counterparts. The dendrogram represents the hierarchical clustering of strains (complete linkage method). Black tiles in the heatmap of the right panel indicate genes whose differential expression is significant (adjusted *p* value < 0.05). Strains belonging to the same Sequence Type (ST) are indicated with shapes.

**Supplementary Fig. 4.**
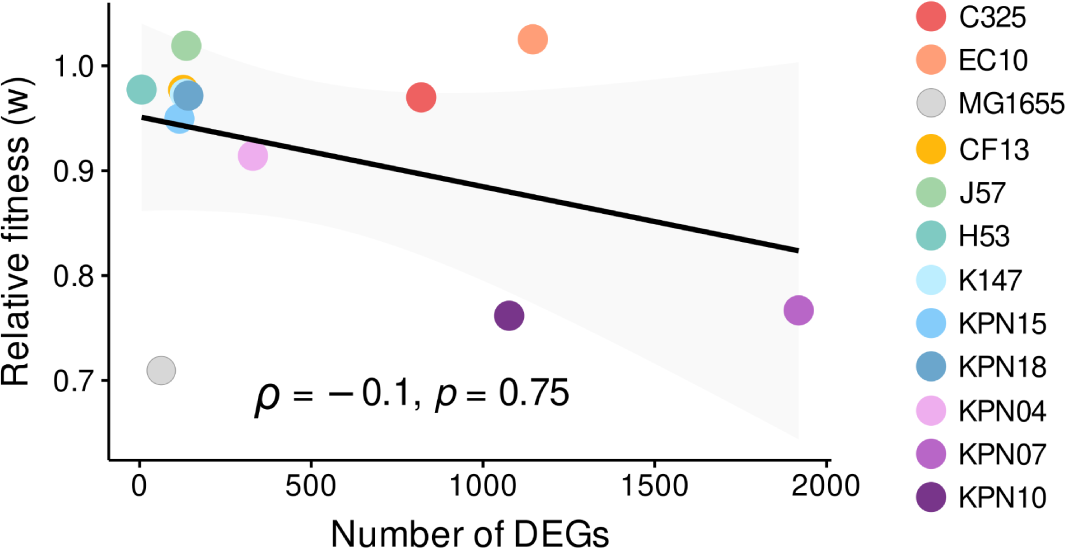
Spearman’s rank correlation between the number of DEGs and the relative fitness (*w*) of pOXA-48-carrying enterobacteria. Relative fitness (*w*) values were previously obtained from competition experiments between pOXA-48-carrying and their respective pOXA-48-free strains ^38–40^. Both variables did not follow a normal distribution (Shapiro test, *P* < 0.05), so Spearman’s rank correlation (*ρ* = -0.1, *P* = 0.75) was used. Gray shading indicates the 95% confidence interval of the regression line (*R*^2^ = 0.14, *F*(1,10) = 1.59, *P* = 0.24; number of DEGs *β* = -6.65 × 10^-5^, *P* = 0.24). Strains are differentiated by colors.

**Supplementary Fig. 5.**
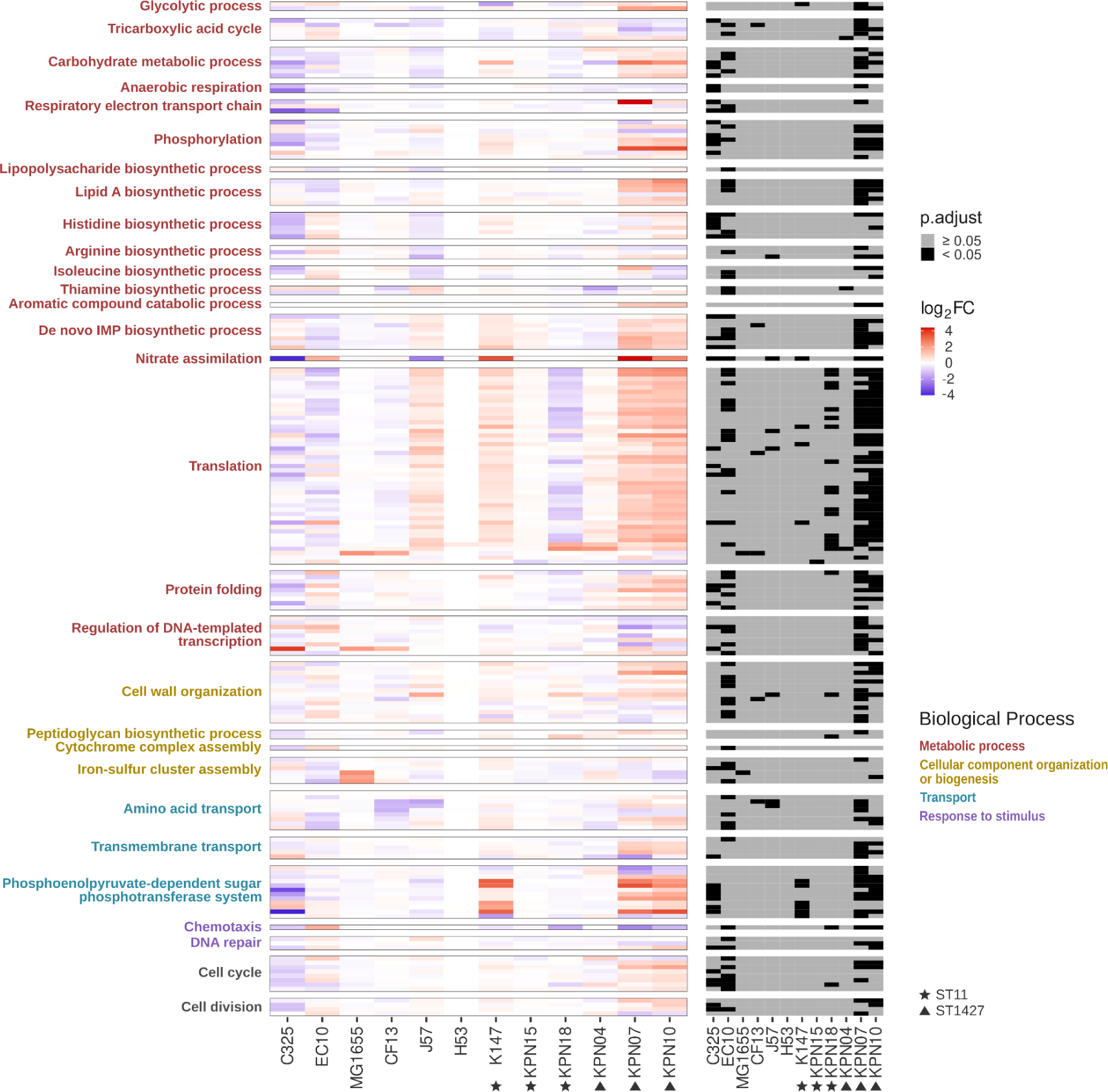
Differential expression of genes from enriched Biological Processes. The heatmap on the left panel represents the differential expression (log_2_FC, calculated by comparing pOXA-48-carrying strains to their pOXA-48-free counterparts) of genes annotated with Biological Processes that were enriched in the GSEA. Genes were matched across strains by RefSeq ID, Gene and Product annotations. Black tiles in the heatmap of the right panel indicate genes whose differential expression is significant (adjusted *p* value < 0.05). Strains belonging to the same Sequence Type (ST) are indicated with shapes.

**Supplementary Fig. 6.**
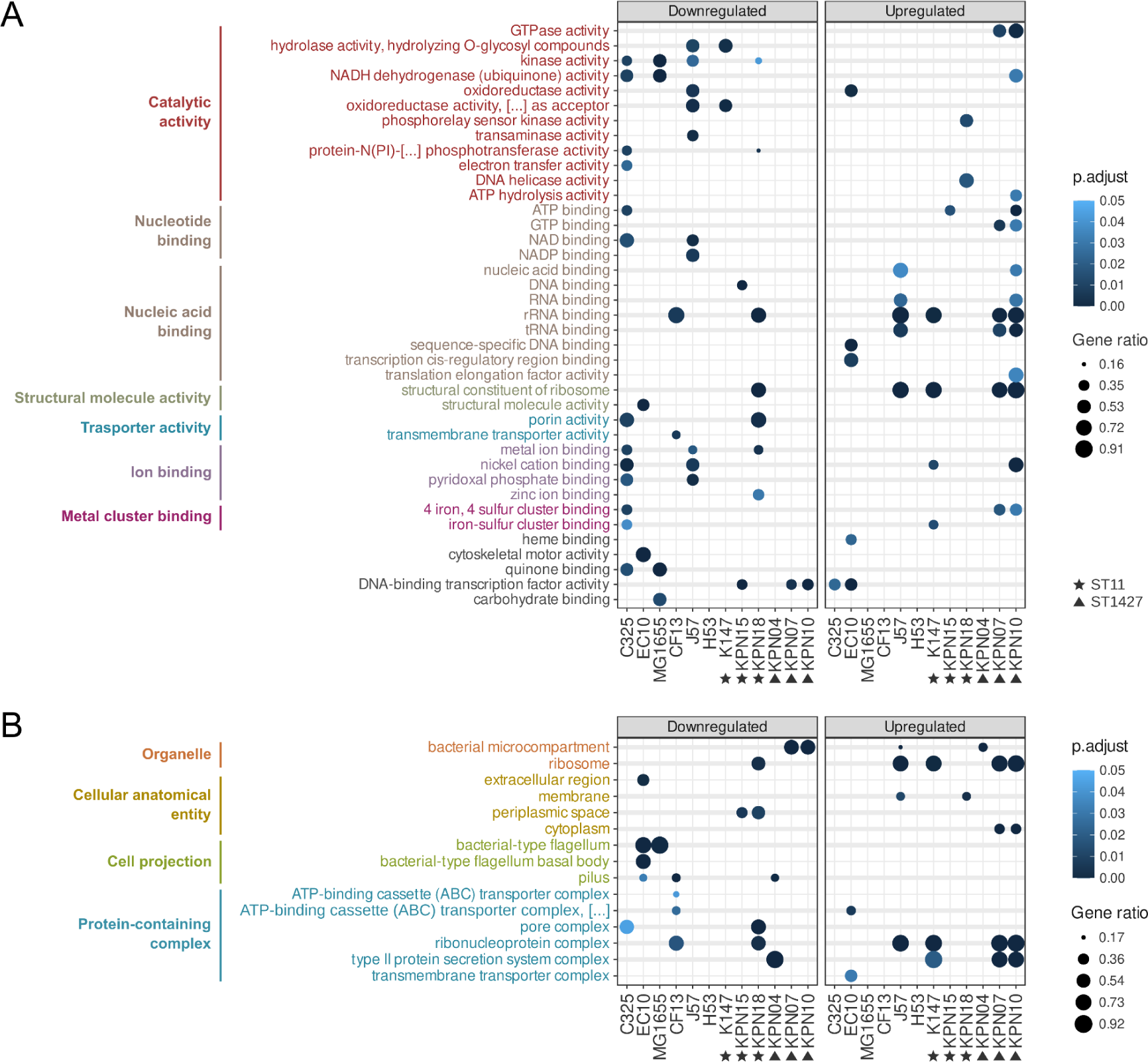
Additional Gene Set Enrichment Analysis (GSEA) results. Molecular Functions (MF) (A) and Cellular Components (CC) (B) enriched in pOXA-48-carrying enterobacteria. Gene Set Enrichment Analysis (GSEA) was performed on the lists of raw DEGs annotated with Gene Ontology terms for MF and CC (see Methods). Downregulated and upregulated enriched MF and CC are separated in two panels, and are represented by circles. Thicker horizontal lines indicate terms enriched in more than one strain. The size of the circles indicates the ratio of the number of enriched genes of a specific MF or CC by the number of total genes annotated with that term. The adjusted *p* value for each enriched term and strain is represented in a color gradient. Strains belonging to the same Sequence Type (ST) are indicated with shapes.

**Supplementary Fig. 7.**
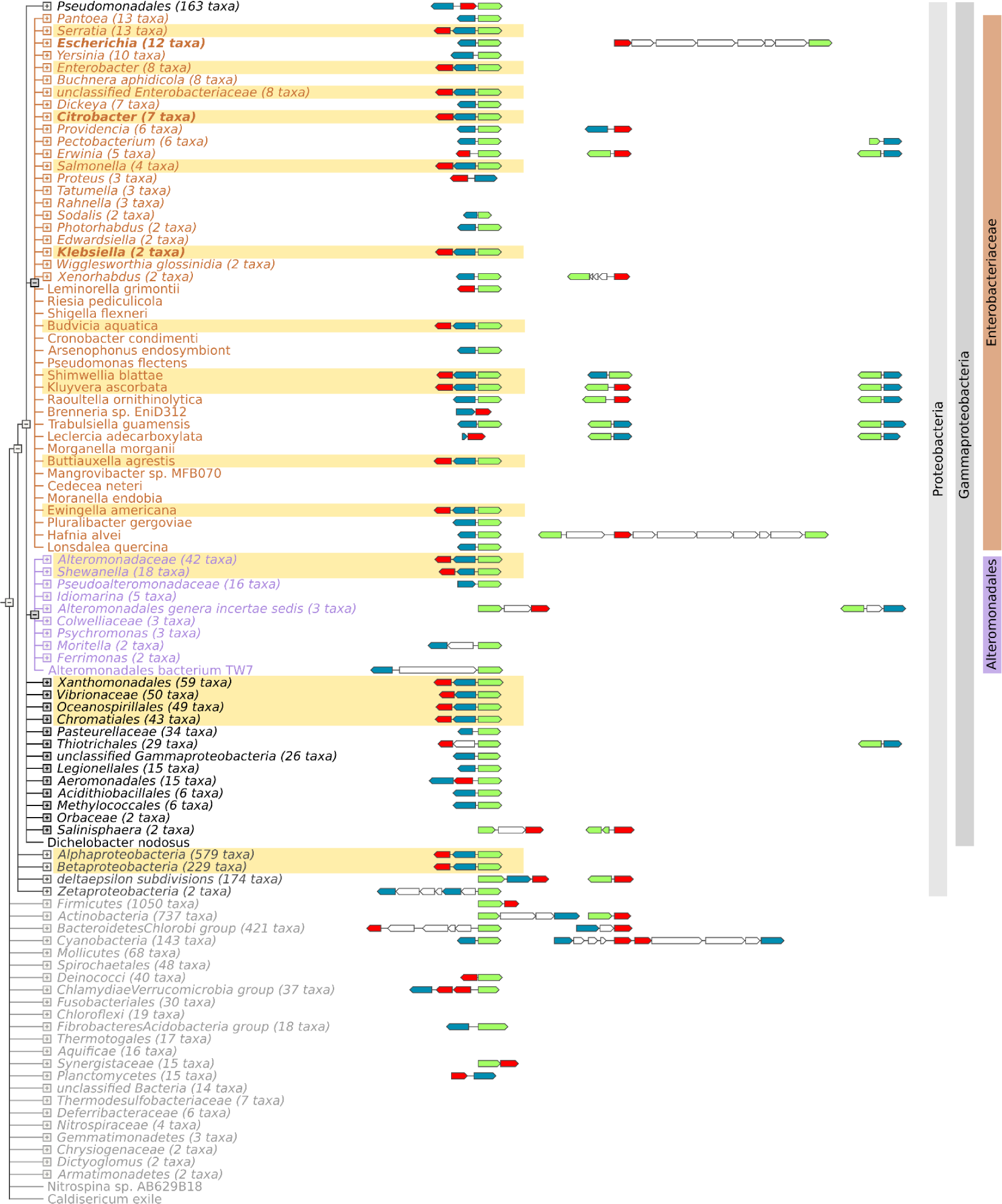
STRING conserved neighborhood view of the *lysR-pfp-ifp* cluster in bacteria. The *lysR*, *pfp* and *ifp* genes are represented as red, blue and green arrows, respectively. The representative neighborhood of the three genes is shown for each bacterial genera/species. Bacterial species or genera encoding the *lysR-pfp-ifp* cluster are highlighted in yellow. Species classification is indicated at the right. The dendrogram does not show phylogenetic distances.

**Supplementary Fig. 8.**
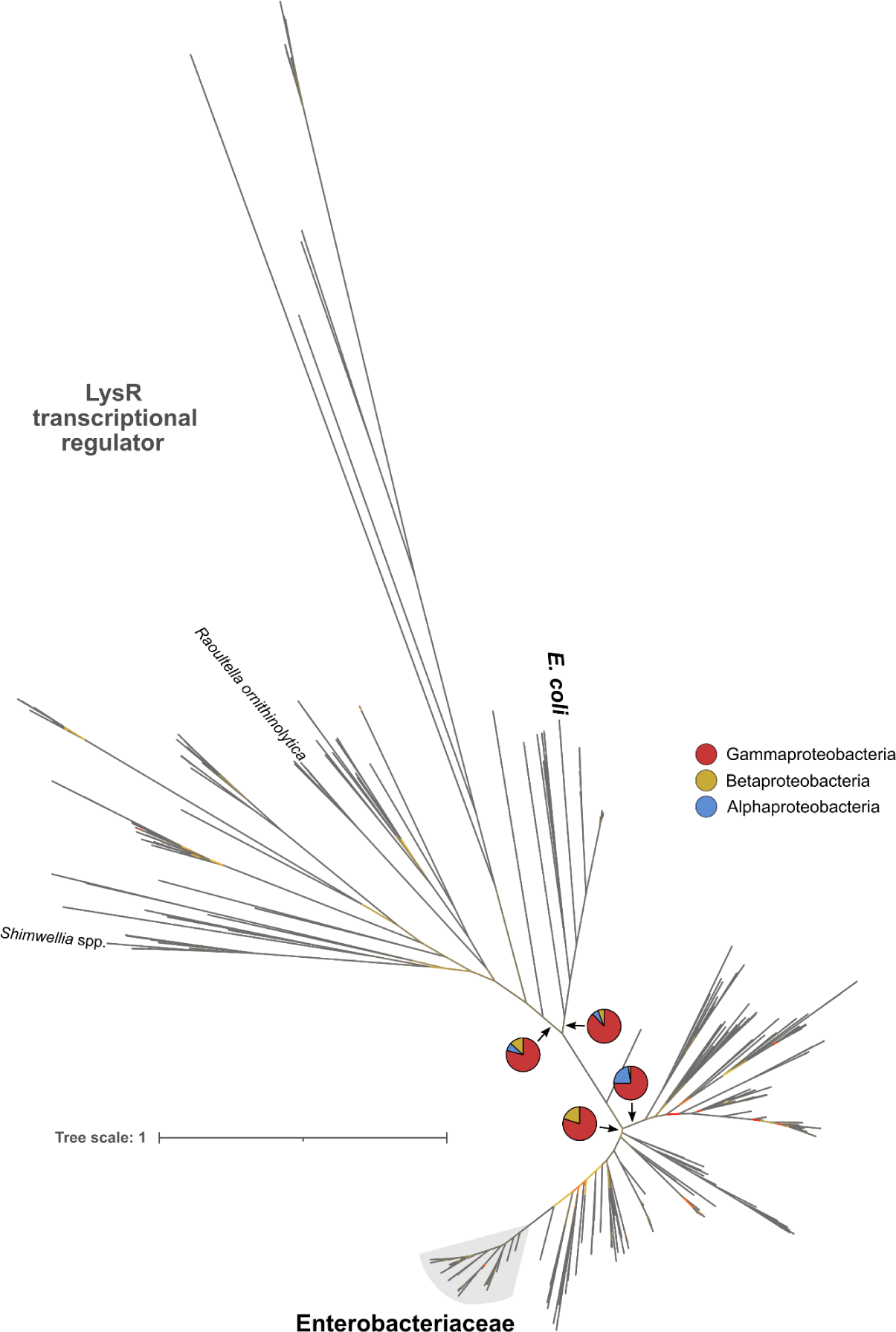
Phylogeny of the LysR transcriptional regulator. Unrooted phylogenetic tree of Proteobacteria (n = 698) constructed from the protein sequences of the LysR that putatively regulates the small chromosomal operon (see Methods). Pie charts show the proportion of Proteobacteria strains belonging to the Gamma-(red), Beta-(yellow) or Alphaproteobacteria (blue) classes included in each of the indicated internal branches. Enterobacteriaceae strains that branch from the same node are shaded in gray; other Enterobacteriaceae species not branching from that node are indicated. Bootstrap values are represented with a color gradient in the branches of the tree: low (≥3), medium and high (≤100) bootstrap values are colored in red, yellow and gray, respectively. Tree scale represents the average number of substitutions per site.

**Supplementary Fig. 9.**
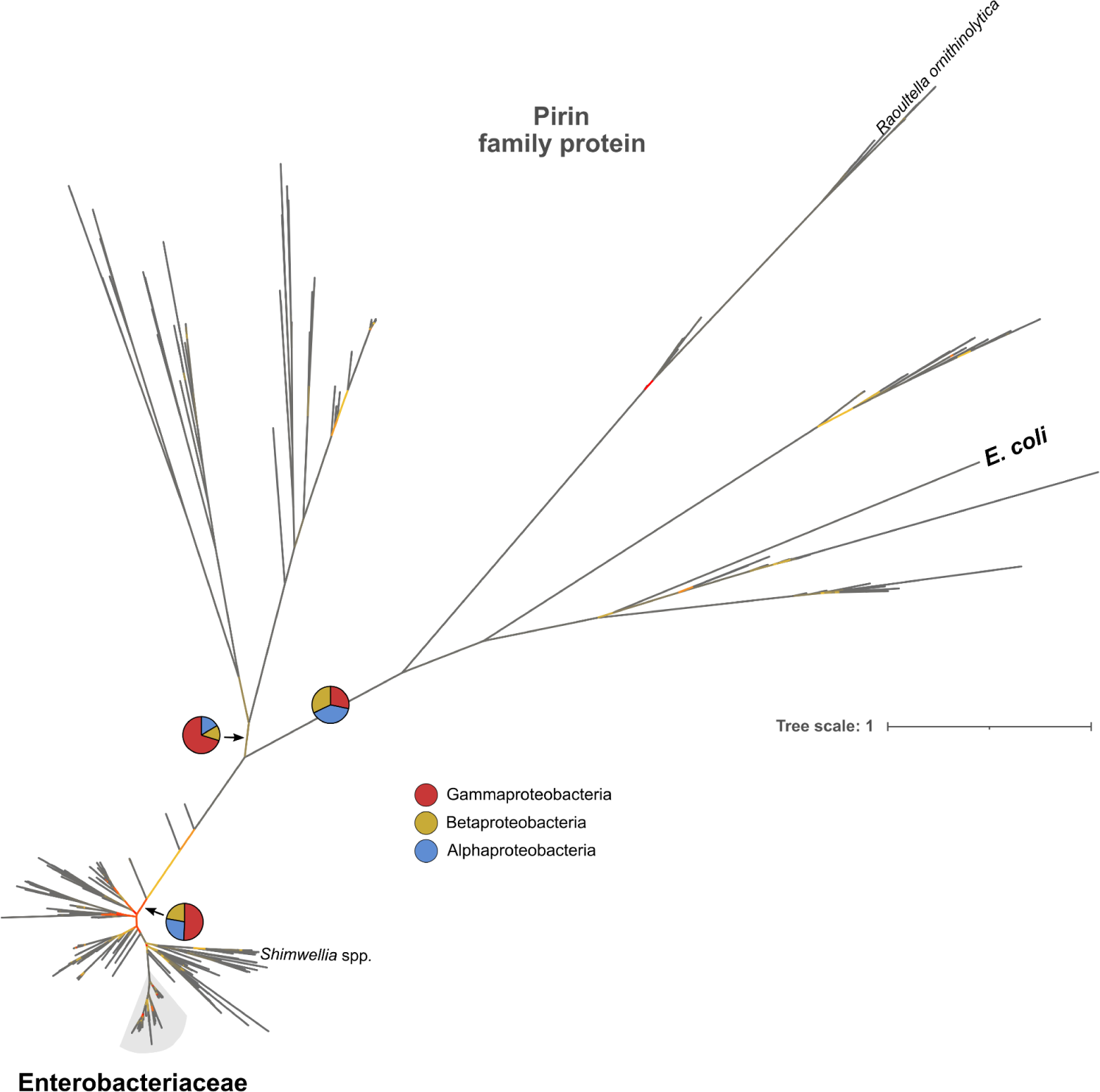
Phylogeny of the pirin family protein. Unrooted phylogenetic tree of Proteobacteria (n = 698) constructed from the protein sequences of the pirin family protein of the small chromosomal operon (see Methods). Pie charts show the proportion of Proteobacteria strains belonging to the Gamma-(red), Beta-(yellow) or Alphaproteobacteria (blue) classes included in each of the indicated internal branches. Enterobacteriaceae strains that branch from the same node are shaded in gray; other Enterobacteriaceae species not branching from that node are indicated. Bootstrap values are represented with a color gradient in the branches of the tree: low (≥3), medium and high (≤100) bootstrap values are colored in red, yellow and gray, respectively. Tree scale represents the average number of substitutions per site.

**Supplementary Fig. 10.**
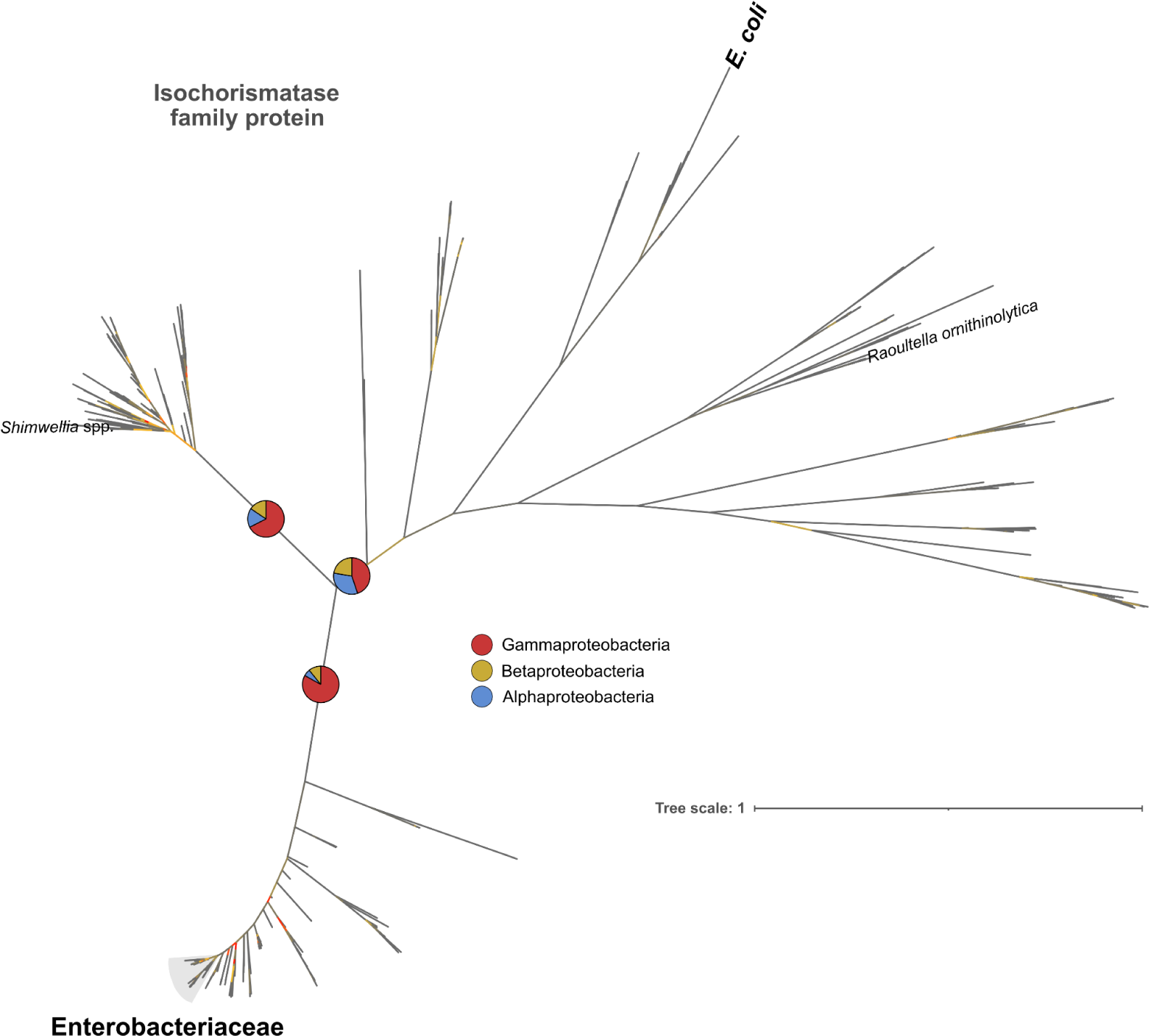
Phylogeny of the isochorismatase family protein. Unrooted phylogenetic tree of Proteobacteria (n = 698) constructed from the protein sequences of the isochorismatase family protein of the small chromosomal operon (see Methods). Pie charts show the proportion of Proteobacteria strains belonging to the Gamma-(red), Beta-(yellow) or Alphaproteobacteria (blue) classes included in each of the indicated internal branches. Enterobacteriaceae strains that branch from the same node are shaded in gray; other Enterobacteriaceae species not branching from that node are indicated. Bootstrap values are represented with a color gradient in the branches of the tree: low (≥1), medium and high (≤100) bootstrap values are colored in red, yellow and gray, respectively. Tree scale represents the average number of substitutions per site.

**Supplementary Fig. 11.**
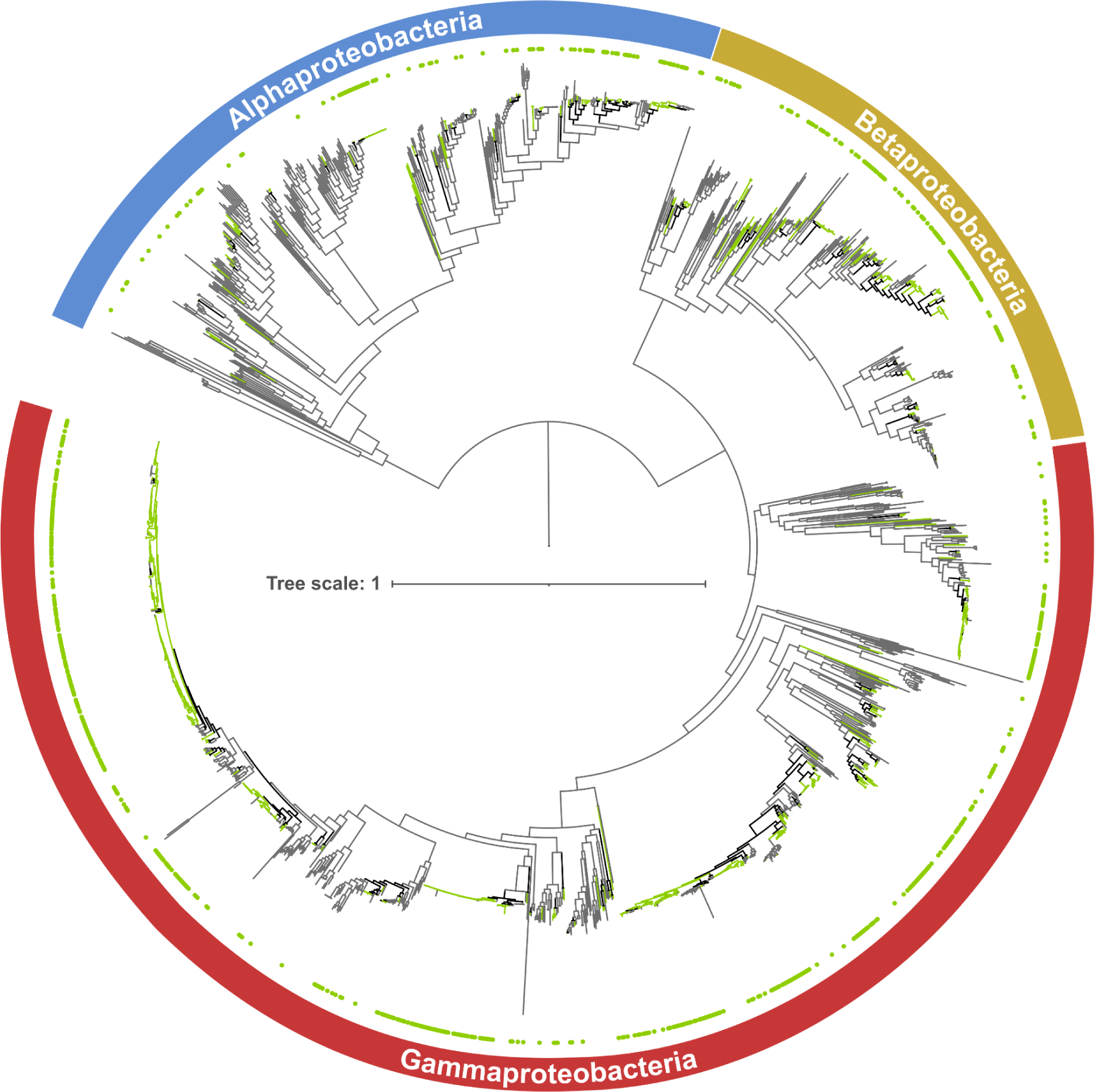
Ancestral character reconstruction of the *lysR-pfp-ifp* cluster. Phylogenetic tree of 1,686 strains of Gamma-, Beta- and Alphaproteobacteria showing the ancestral character reconstruction (ACR) of the *lysR-pfp-ifp* cluster, rooted at the Epsilonproteobacteria outgroup (removed from the tree). The tree was constructed from the concatenated protein sequences of 128 conserved bacterial single-copy genes (see Methods). Green dots at the tips of the tree and green nodes/branches indicate presence of the *lysR-pfp-ifp* cluster. Gray and black nodes/branches indicate absence or an undefined state (after ACR) of the *lysR-pfp-ifp* cluster, respectively. See Supplementary Fig. 19 (unpruned tree) for bootstrap values. Tree scale represents the average number of substitutions per site.

**Supplementary Fig. 12.**
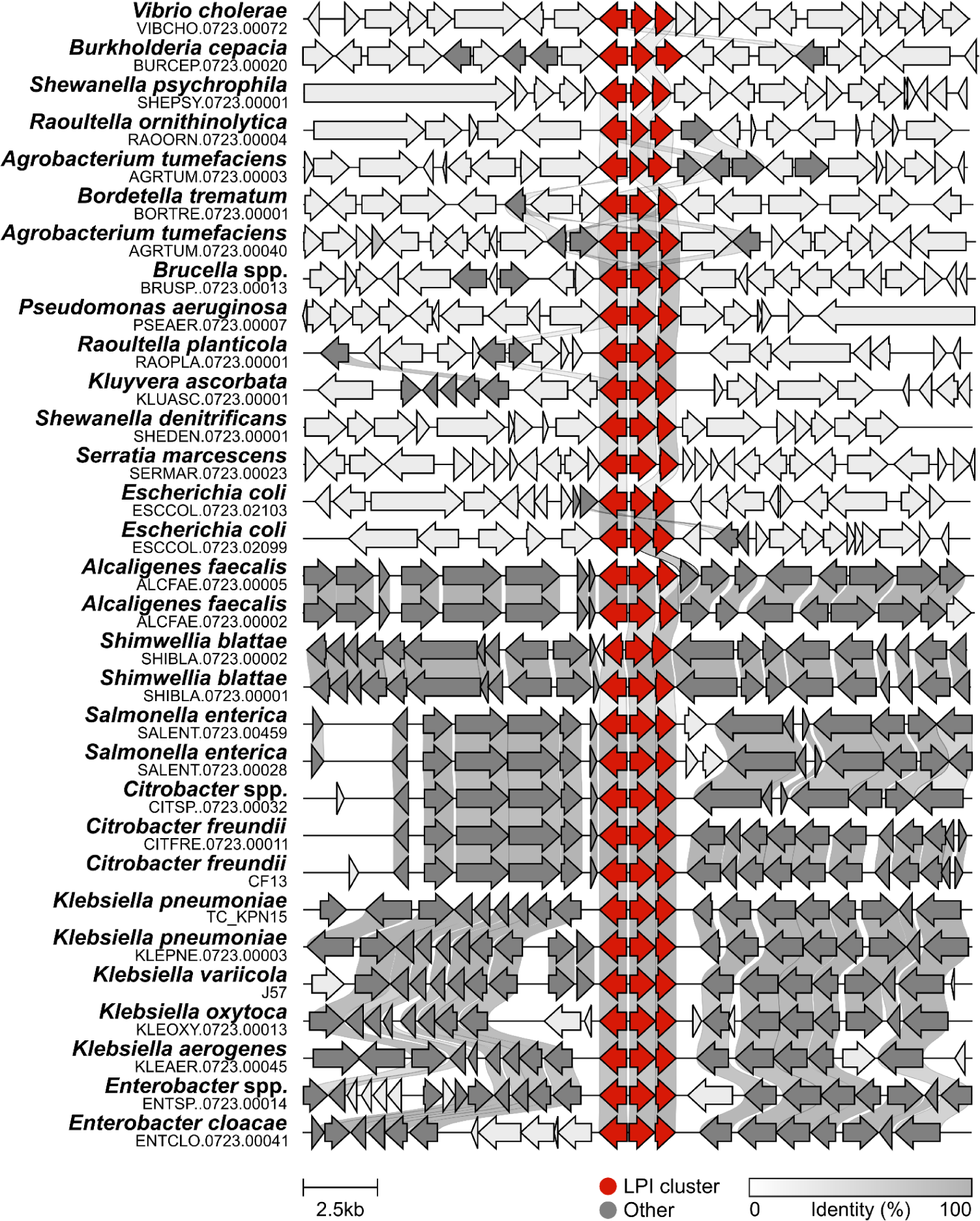
Genomic neighborhood of the *lysR-pfp-ifp* cluster. A representative subset of Proteobacteria strains was selected (see Methods). Genes are color-coded as indicated at the bottom. Homologous genes between strains and their percentage of sequence identity are indicated with a gray shading.

**Supplementary Fig. 13.**
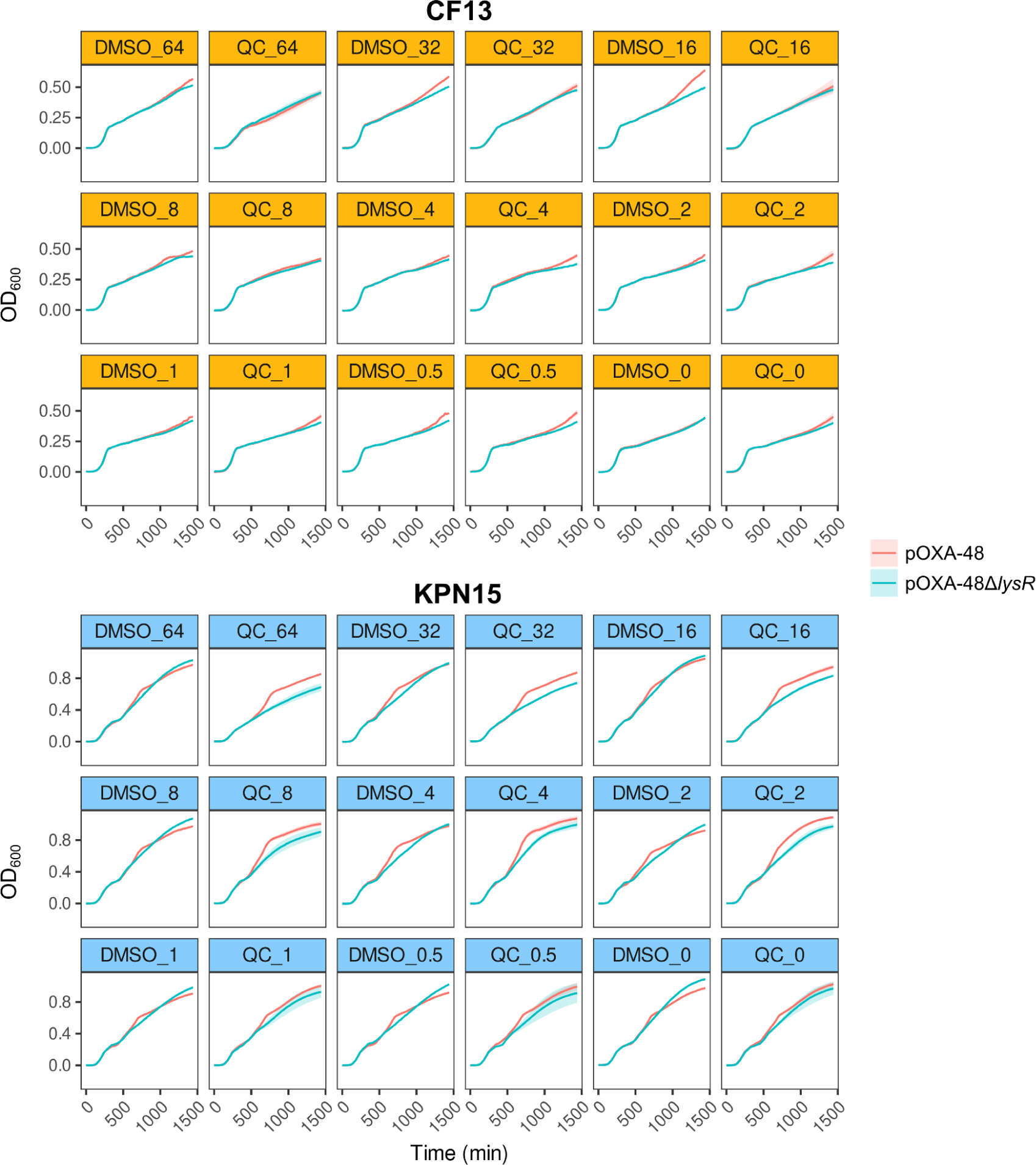
Growth curves of clinical enterobacteria grown under different concentrations of quercetin (QC). OD_600_ measured every 10 minutes for 24 hours for strains CF13 and KPN15 carrying pOXA-48 or pOXA-48Δ*lysR.* Strains were grown under concentrations ranging from 64 to 0.5 µg/mL of DMSO (as control, 1 replicate per concentration and strain) or DMSO+QC (2 replicates per concentration and strain). Shading indicates the standard error of the mean.

**Supplementary Fig. 14.**
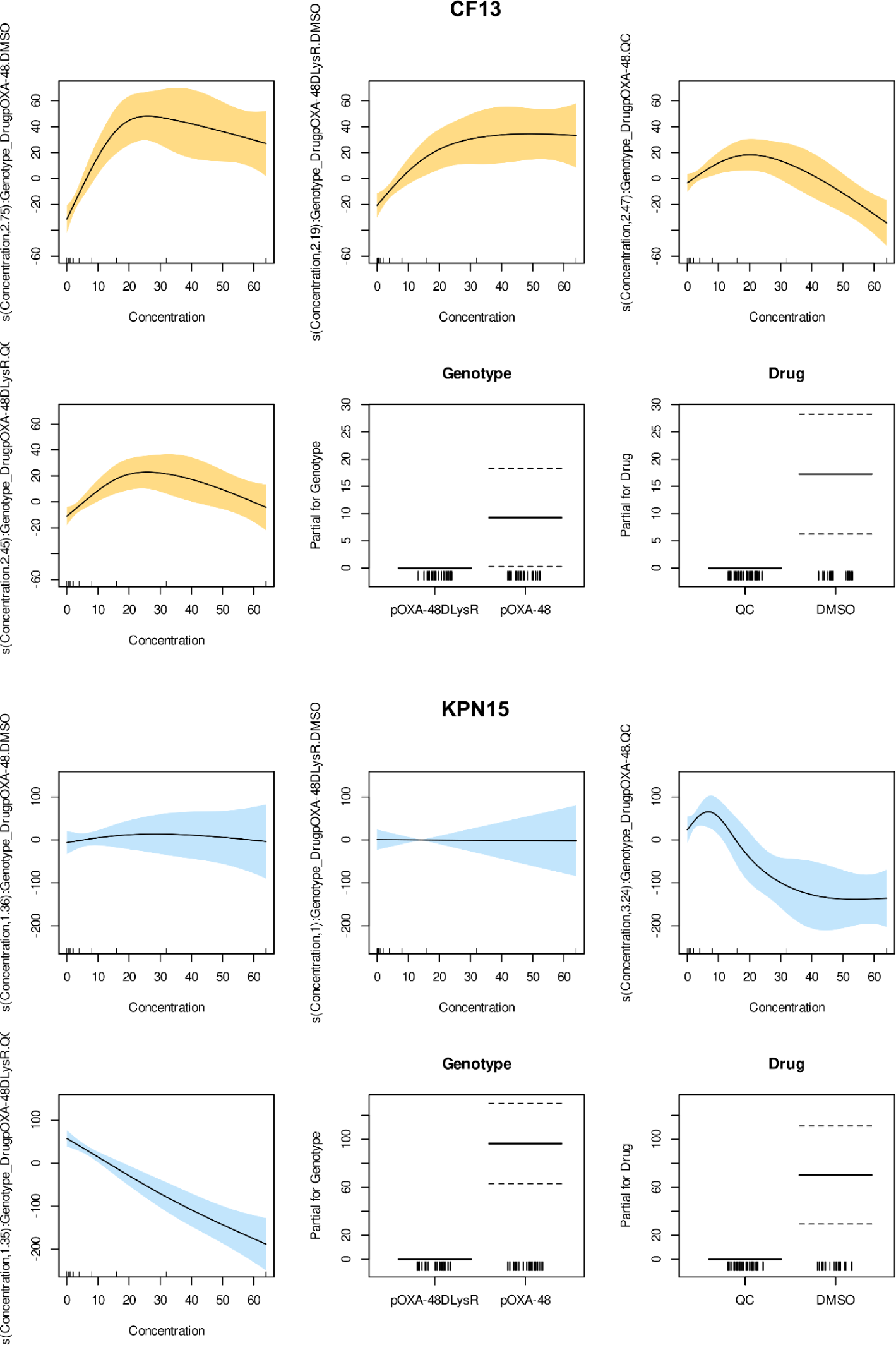
Generalized Additive Model (GAM) results. The GAMs are fitted to model bacterial growth (AUC of growth curves, Supplementary Fig. 13) as a function of the concentration of the drugs (DMSO and DMSO+QC), for CF13 (top) and KPN15 (bottom) separately. Four smooth functions are fitted for each genotype (pOXA-48 or pOXA-48Δ*lysR*) and drug (DMSO and DMSO+QC) interaction combinations, with the shaded area indicating the 95% confidence interval. Interaction effects of genotype and drug are incorporated into the model. Genotype pOXA-48Δ*lysR* and drug DMSO+QC are set as reference levels. Rug plots along the x-axis indicate the observed data points. Models fitted using the restricted maximum likelihood (REML) method (see Methods).

**Supplementary Fig. 15.**
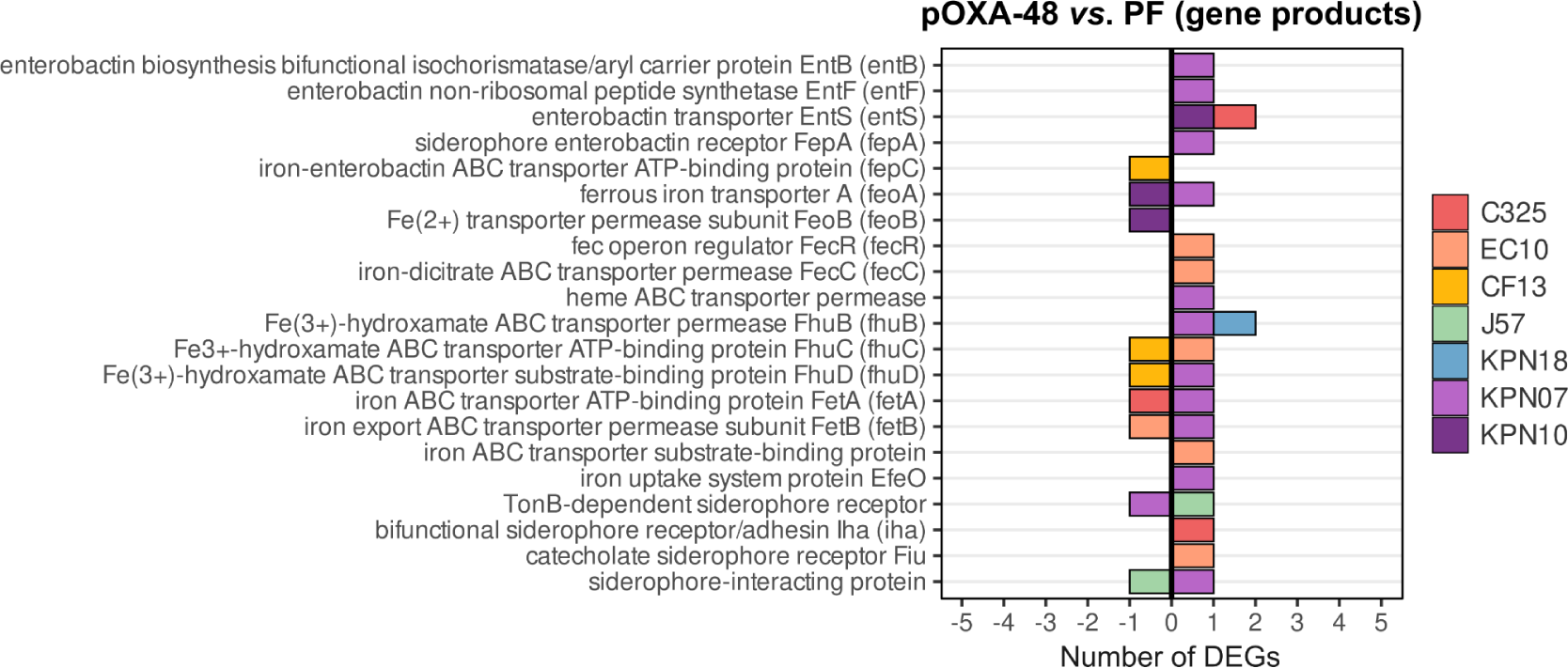
Number of significant DEGs involved in iron transport. DE was analyzed by comparing pOXA-48*-*carrying against pOXA-48-free strains (first RNA-Seq dataset). Values >0 represent the count of upregulated genes, and values <0 are downregulated genes.

**Supplementary Fig. 16.**
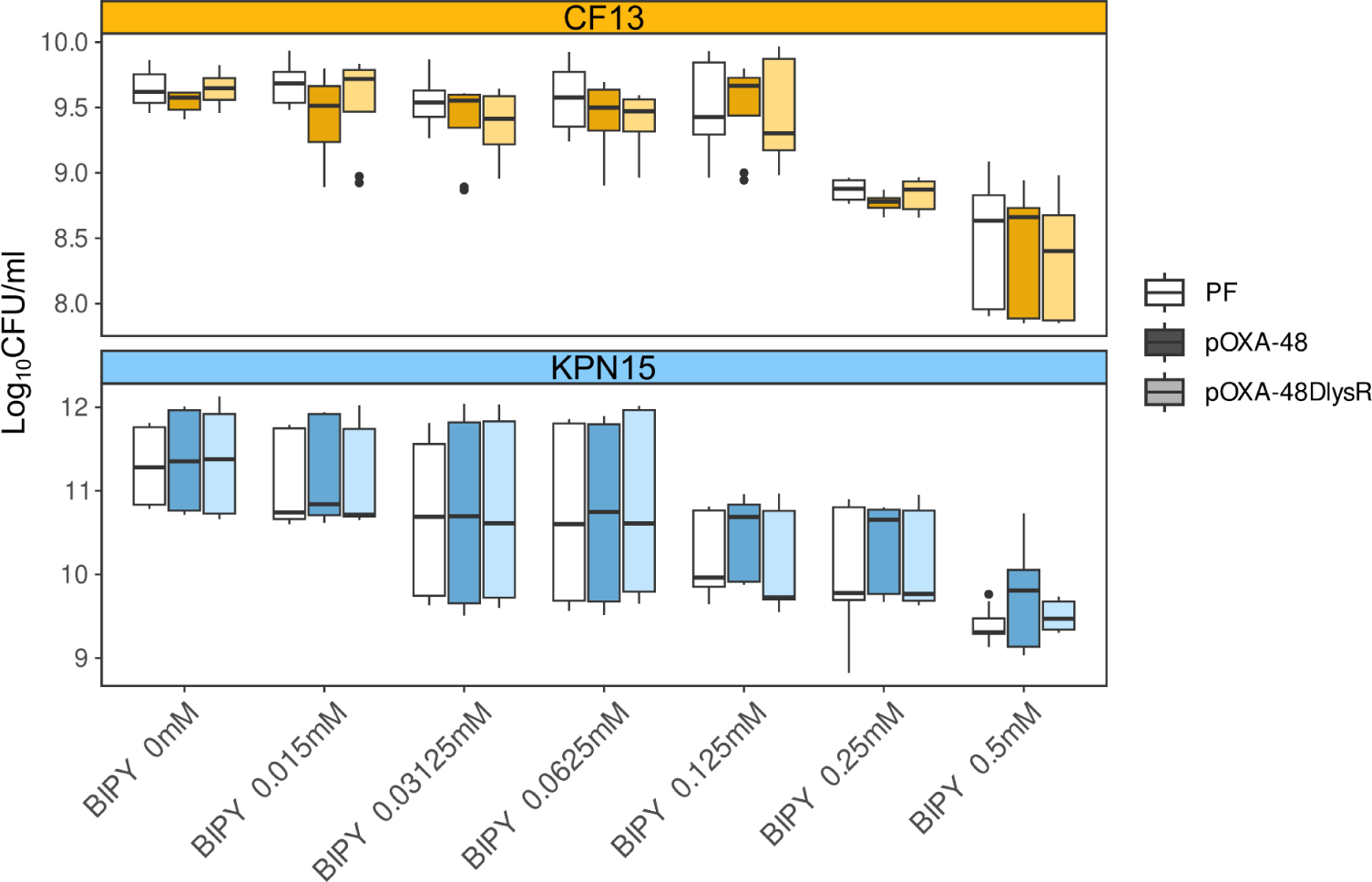
Growth of clinical enterobacteria under increasing concentrations of the iron chelator bipyridine. Colony forming units (CFU) per mL of strains CF13 (top figure) and KPN15 (bottom figure) not carrying or carrying pOXA-48 or pOXA-48Δ*lysR* grown under increasing concentrations of the iron chelator bipyridine. Median CFU/mL values are indicated as horizontal lines inside the boxes, the upper and lower box hinges correspond to the 25th and 75th percentiles and the whiskers extend to observations within 1.5x the interquartile range.

**Supplementary Fig. 17.**
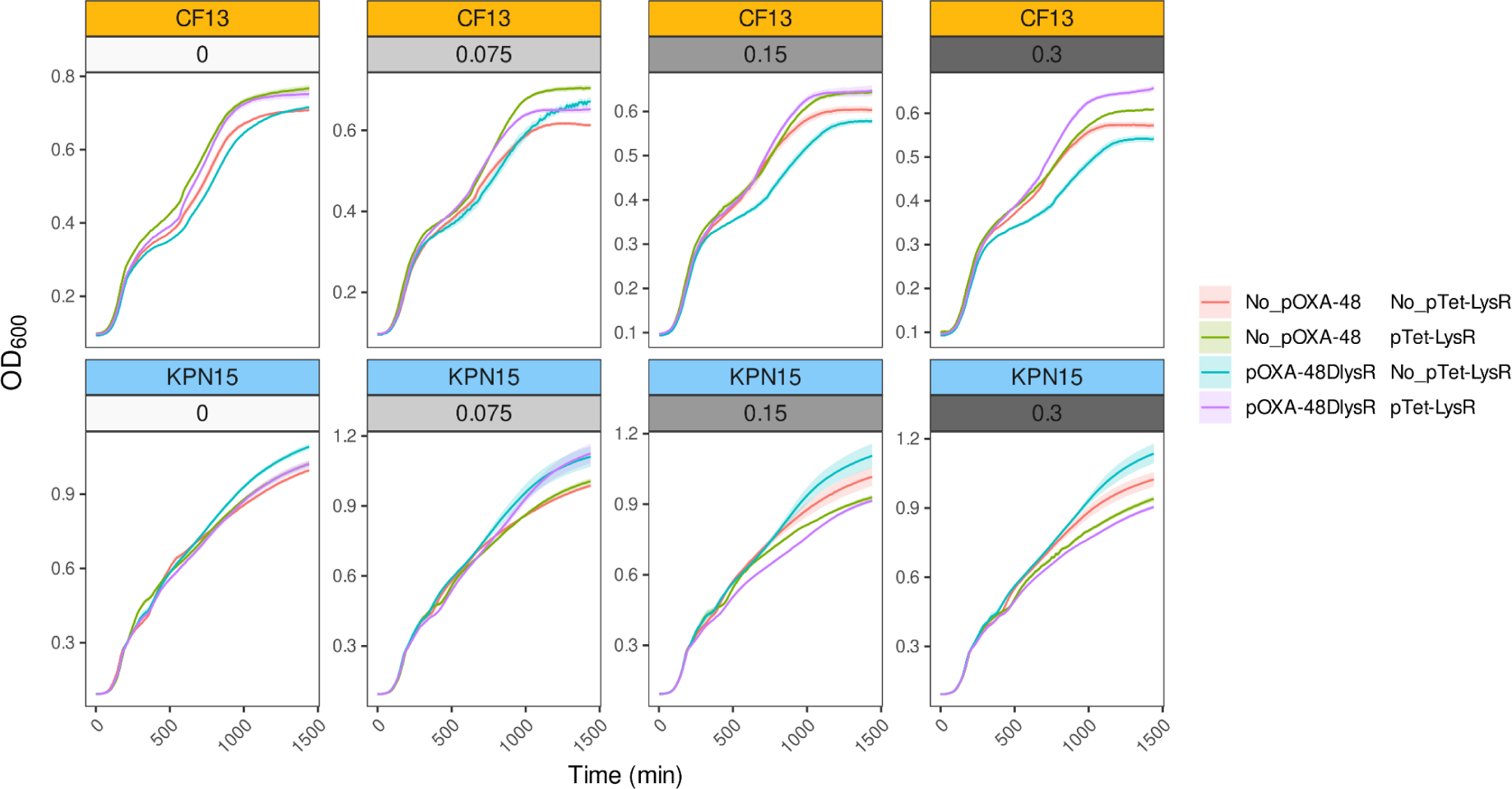
Growth curves of strains overexpressing LysR_pOXA-48_ at different concentrations of inducer. Effect of the overexpression of LysR_pOXA-48_ on the growth of CF13 and KPN15. Curves performed for plasmid-free and pOXA-48Δ*lysR*-carrying CF13 and KPN15, with expression of LysR_pOXA-48_ in *trans* from the pTet-LysR. Experiment performed with increasing concentrations of inducer (aTc). Shading indicates the standard error of the mean.

**Supplementary Fig. 18.**
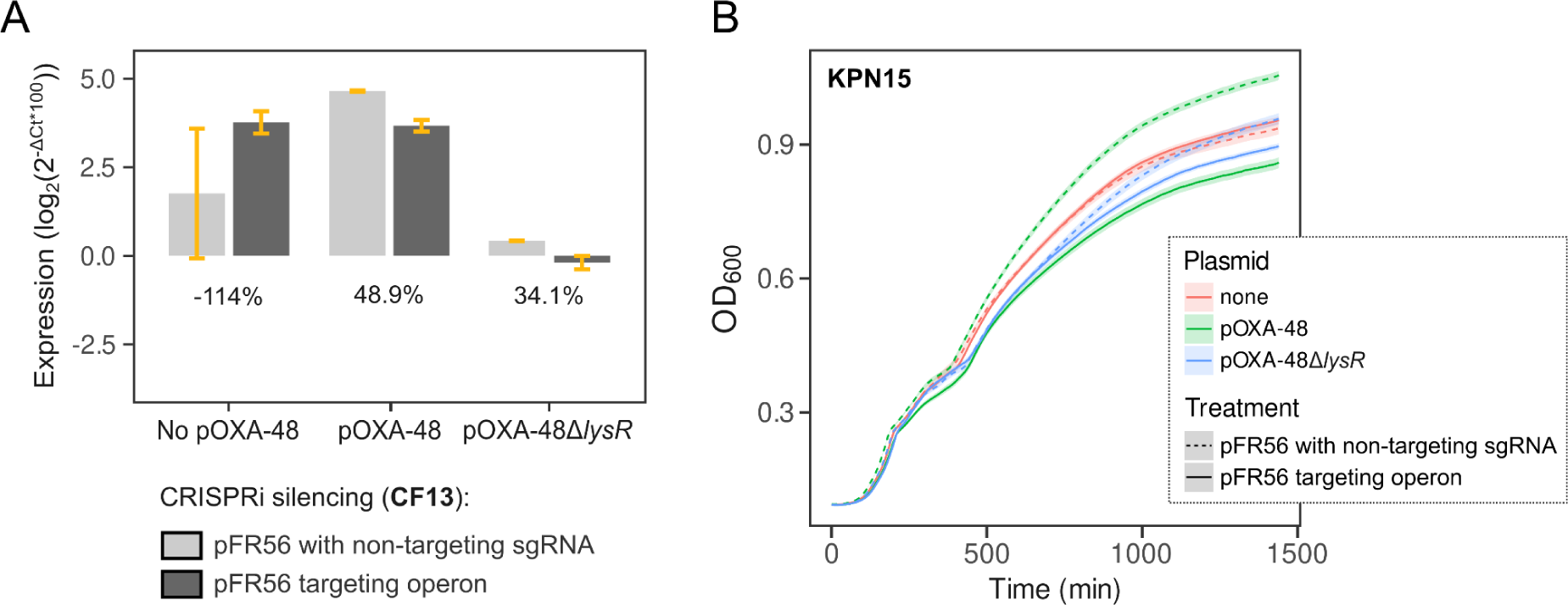
Repression of the *pfp-ifp* operon using CRISPRi. (A) RT-qPCR results for the level of repression of the *pfp-ifp* operon using CRISPRi in CF13. Levels of expression of the operon are calculated using RT-qPCR results for *pfp* as proxy for the whole operon and are normalized to the endogenous control *rpoB*. Percentages indicate the percentage of repression of the operon comparing the expression levels of cells carrying the non-targeting version of pFR56apm and cells carrying pFR56apm targeting the operon. (B) Growth curves (optical density at 600 nm wavelength) of strain KPN15 with CRISPRi silencing of the *pfp-ifp* operon, used to calculate AUC values for Fig. 7C. OD_600_ was measured every 10 minutes during 24 hours for eight replicates per group. Error bars (A) and shading (B) indicate the standard error of the mean.

**Supplementary Fig. 19.**
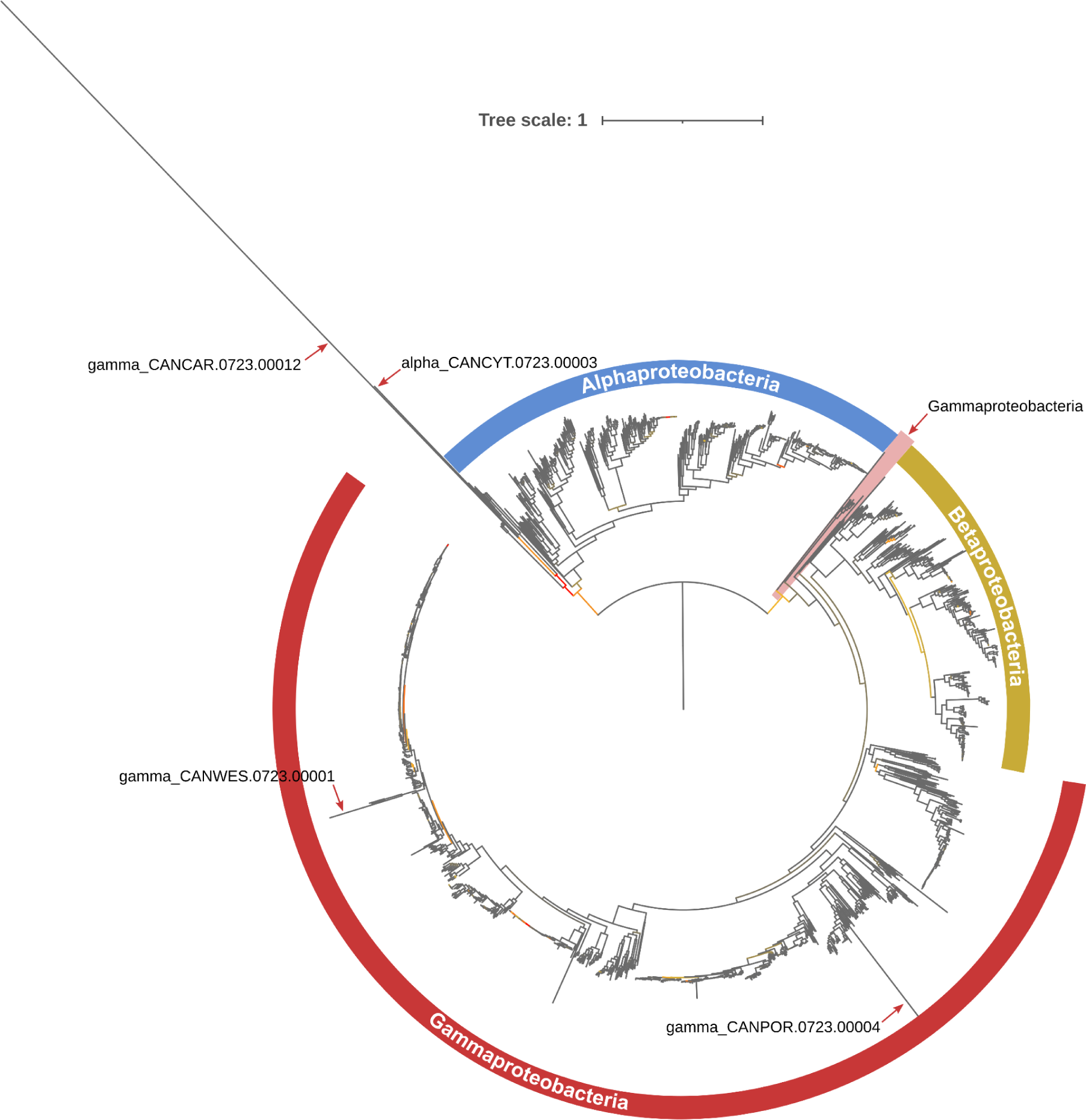
Unpruned phylogenetic tree of Proteobacteria. Phylogenetic tree of 1,704 strains of Gamma-, Beta- and Alphaproteobacteria, rooted at the Epsilonproteobacteria outgroup (removed from the tree). The tree was constructed from the concatenated protein sequences of 128 conserved bacterial single-copy genes (see Methods). The four outlier taxa identified by TreeShrink (Candidatus *Cytomitobacter primus* [GCF_951805245.1], Candidatus *Carsonella ruddii* [GCF_016593055.1], Candidatus *Westeberhardia cardiocondylae* [GCF_001242845.1], Candidatus *Portiera aleyrodidarum* BT-B-HRs [GCF_000300075.1]) and the clade of Gammaproteobacteria branching outside of the Gammaproteobacteria clade (*Sulfurifustis variabilis* [GCF_002355415.1], Candidatus *Ruthia magnifica* str. Cm [GCF_000015105.1], *Thiosulfatimonas sediminis* [GCF_011398355.1], *Hydrogenovibrio crunogenus* [GCF_004786015.1], *Ignatzschineria rhizosphaerae* [GCF_022655595.1], *Francisella frigiditurris* [GCF_001880225.1], *Francisella* sp. Scap27 [GCF_013394105.1], *Francisella adeliensis* [GCF_012224465.1], *Francisella halioticida* [GCF_002211785.1], *Francisella persica* ATCC VR-331 [GCF_001653955.1], *Francisella opportunistica* [GCF_003347095.1], *Francisella philomiragia* [GCF_000833315.1], *Francisella noatunensis* subsp. *noatunensis* FSC774 [GCF_014844275.1], Candidatus *Comchoanobacter bicostacola* [GCF_024172705.1]) are indicated. These were pruned from the tree prior to ACR (Supplementary Fig. 11). Bootstrap values are represented with a color gradient in the branches of the tree: low (≥8), medium and high (≤100) bootstrap values are colored in red, yellow and gray, respectively. Tree scale represents the average number of substitutions per site.

**Supplementary Fig. 20.**
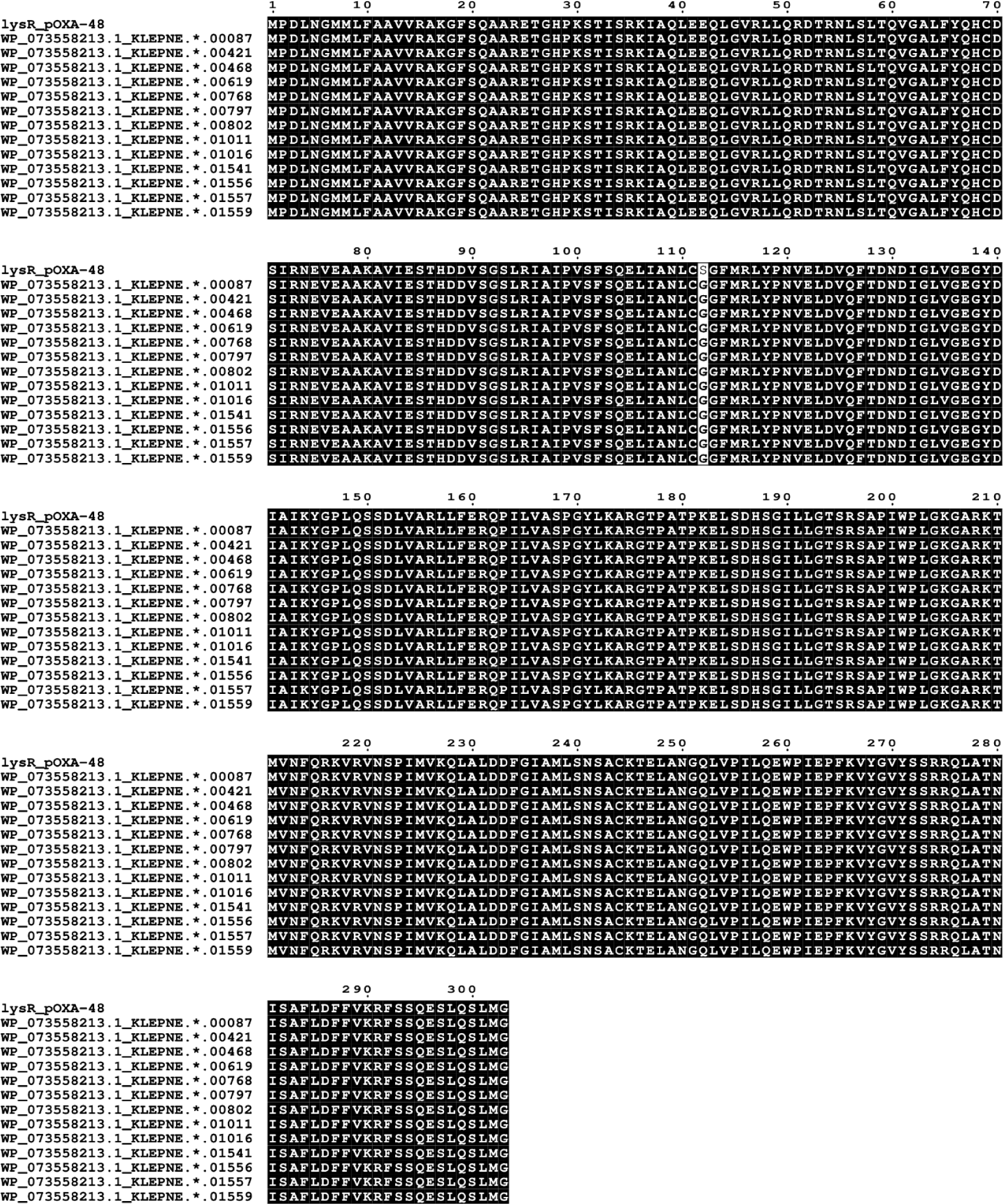
MAFFT alignment of plasmid-encoded LysR transcriptional regulators. The protein sequences of 13 plasmid-encoded LysRs that share less than 100% (and >96%) identity with LysR_pOXA-48_ are aligned to LysR_pOXA-48_ (top of alignment). Matching and mismatching amino acids are highlighted in black and white, respectively. Of the 13 LysRs, eight are encoded in IncL/Ms (pOXA-48 plasmid variants): WP_073558213.1 in KLEPNE.0723.00087, WP_073558213.1 in KLEPNE.0723.00421, WP_073558213.1 in KLEPNE.0723.00468, WP_073558213.1 in KLEPNE.0723.00619, WP_073558213.1 in KLEPNE.0723.00768, WP_073558213.1 in KLEPNE.0723.01011, WP_073558213.1 in KLEPNE.0723.01016 and WP_073558213.1 in KLEPNE.0723.01556.

